# GRAMD1C regulates autophagy initiation and mitochondrial bioenergetics through ER-mitochondria cholesterol transport

**DOI:** 10.1101/2021.08.04.455032

**Authors:** Matthew Yoke Wui Ng, Chara Charsou, Ana Lapao, Sakshi Singh, Laura Trachsel-Moncho, Sigve Nakken, Michael J. Munson, Anne Simonsen

## Abstract

During autophagy, cytosolic cargo is sequestered into double-membrane vesicles called autophagosomes. The origin and identity of the membranes that form the autophagosome remain to be fully characterized. Here, we investigated the role of cholesterol in starvation- induced autophagy and identify a role for the ER-localized cholesterol transport protein GRAMD1C in the regulation of autophagy and mitochondrial function. We demonstrate that cholesterol depletion leads to a rapid induction of autophagy, possibly caused by a corresponding increased abundance of curved autophagy membranes. We further show that GRAMD1C is a negative regulator of starvation-induced autophagy. Similar to its yeast orthologue, GRAMD1C is recruited to mitochondria through its GRAM domain. Additionally, we find that GRAMD1C depletion leads to increased mitochondrial cholesterol accumulation and mitochondrial oxidative phosphorylation. Finally, we demonstrate that expression of GRAM family genes is linked to clear cell renal carcinoma survival, highlighting the pathophysiological relevance of cholesterol transport proteins.

## Introduction

Macroautophagy (referred henceforth as autophagy) involves the *de novo* formation of membranes that sequester cytoplasmic cargo into double-membrane autophagosomes, which subsequently fuse with lysosomes, leading to cargo degradation and recycling of the resulting macromolecules to obtain homeostasis during periods of starvation and cellular stress. Autophagosome biogenesis is initiated through the recruitment of the ULK1 kinase complex (including ULK1, ATG101, ATG13, FIP200) and the class III phosphatidylinositol 3-kinase complex 1 (PIK3C3-C1, consisting of BECN1, ATG14L1, PIK3C3, PIK3R4) to endoplasmic reticulum (ER) associated sites, from where newly formed autophagosomes emanate. The autophagosome membrane is largely devoid of transmembrane proteins^1^ and its formation is therefore thought to be regulated by membrane associated proteins, as well as its lipid composition and lipid distribution^2^. Previous studies have shown that autophagosomes are enriched in unsaturated fatty acids^3^ and that these are necessary for autophagosome formation^4, 5^. These observations suggest that decreased membrane order is favorable towards autophagy initiation, possibly by generating flexible, highly curved membranes known to be required for autophagosome formation^6^. In addition to phospholipids, cholesterol is a crucial component of mammalian membranes and its abundance is also a determinant of membrane order and fluidity^7, 8^. It is noteworthy that freeze fracture electron microscopy analysis revealed early autophagosomal structures to be cholesterol poor^9^, suggesting that cholesterol poor membranes are the principal source of membranes during autophagosome biogenesis. In agreement with this, cholesterol depletion with methyl-β cyclodextrin (MBCD) and statins have been reported to promote LC3 lipidation and its turnover^10–15^. A few studies have however found that high cholesterol levels promote autophagy^16, 17^. The majority of these studies involved long-term cholesterol manipulation that can lead to metabolic rewiring, and changes in transcriptional and signaling pathways^18^ and might therefore not reflect the direct influence of cholesterol in autophagy.

Intracellular cholesterol levels are maintained through a combination of new synthesis and extracellular cholesterol uptake. In addition to the low-density lipoprotein receptor (LDLR) pathway, several cholesterol transport proteins have been shown to mediate import and intracellular cholesterol transport directly from the plasma membrane (PM)^19, 20^. An example of such cholesterol transport proteins is the GRAM family (consisting of GRAMD1A, GRAMD1B, GRAMD1C, GRAMD2 and GRAMD3, also known as Aster Proteins), named after the PH-like GRAM domain in their N-terminal region. Among the five members, only GRAMD1A, GRAMD1B and GRAMD1C (herein collectively referred to as GRAMD1s) contain a sterol binding VASt domain, allowing them to facilitate PM to ER cholesterol import^21–23^. Loss of GRAMD1s lead to accumulation of cholesterol on the plasma membrane^21^, and mouse macrophages lacking GRAMD1A and GRAMD1B displayed increased cholesterol accumulation in the PM and upregulated expression of SREBP2 target genes, indicative of decreased ER cholesterol^23^. Due to a limited number of studies on the GRAM family proteins, their biological importance is still not fully understood. A recent study reported that GRAMD1A activity is required for autophagy initiation^24^, suggesting that cholesterol transport proteins can facilitate autophagy through regulation of cholesterol movement. GRAM family proteins were shown to form heterocomplexes in a manner that is dependent on their C-terminal amphipathic helix region^21^, indicating that other GRAM proteins may also regulate autophagy.

Here, we show that cholesterol and the ER-anchored cholesterol transport protein GRAMD1C negatively regulates autophagy initiation. We show that GRAMD1C associates with mitochondria through its GRAM domain and that its depletion leas to accumulation of mitochondrial cholesterol and increased mitochondrial respiration. Finally, we show that members of the GRAM family of cholesterol transport proteins are involved in clear cell renal carcinoma (ccRCC) survival. The GRAMs therefore represent potential regulators of the autophagy pathway and ccRCC.

## Results

### Cholesterol depletion promotes autophagy initiation

Previous *in vivo* and *in vitro* studies have indicated that cholesterol depletion promotes autophagy^13, 25–27^, but most of these studies involve cells being depleted of cholesterol for extended time periods where changes in autophagy can be caused by metabolic and transcriptional responses, thus indirectly activating autophagy. To investigate a more direct role of cholesterol on membrane remodeling during autophagy, we analyzed the short-term effects on starvation-induced autophagy after cholesterol depletion using MBCD, which rapidly removes cholesterol from cellular membranes^28^. U2OS cells were treated with MBCD for 1 hr in control (DMEM) or starvation (EBSS) medium in the presence or absence of the lysosomal V-ATPase inhibitor Bafilomycin A1 (BafA1), followed by immunoblotting for the autophagosome marker LC3B and p62, to determine autophagic flux. Cholesterol depletion caused a 3-fold increase in LC3B lipidation (LC3B-II) under basal conditions, compared to control cells, which was further enhanced in starved cells depleted of cholesterol (Figure 1a-b). In line with this, cholesterol depletion increased the formation of endogenous LC3B puncta both in fed and starved cells, as analyzed by immunofluorescence microscopy (Figure 1c-d). In both cases, the starvation-induced autophagic flux was higher in MBCD treated cells, suggesting that cholesterol and amino acid depletion can modulate autophagy synergistically. Similarly, long-term cholesterol depletion using atorvastatin (ATV) for 48 hrs, an HMG-CoA reductase inhibitor, caused a similar increase in starvation-induced autophagy (Figure 1e-f), while p62 turnover was not significantly altered with MBCD or ATV treatment (Supplementary figure 1a-b).

**Figure 1.**
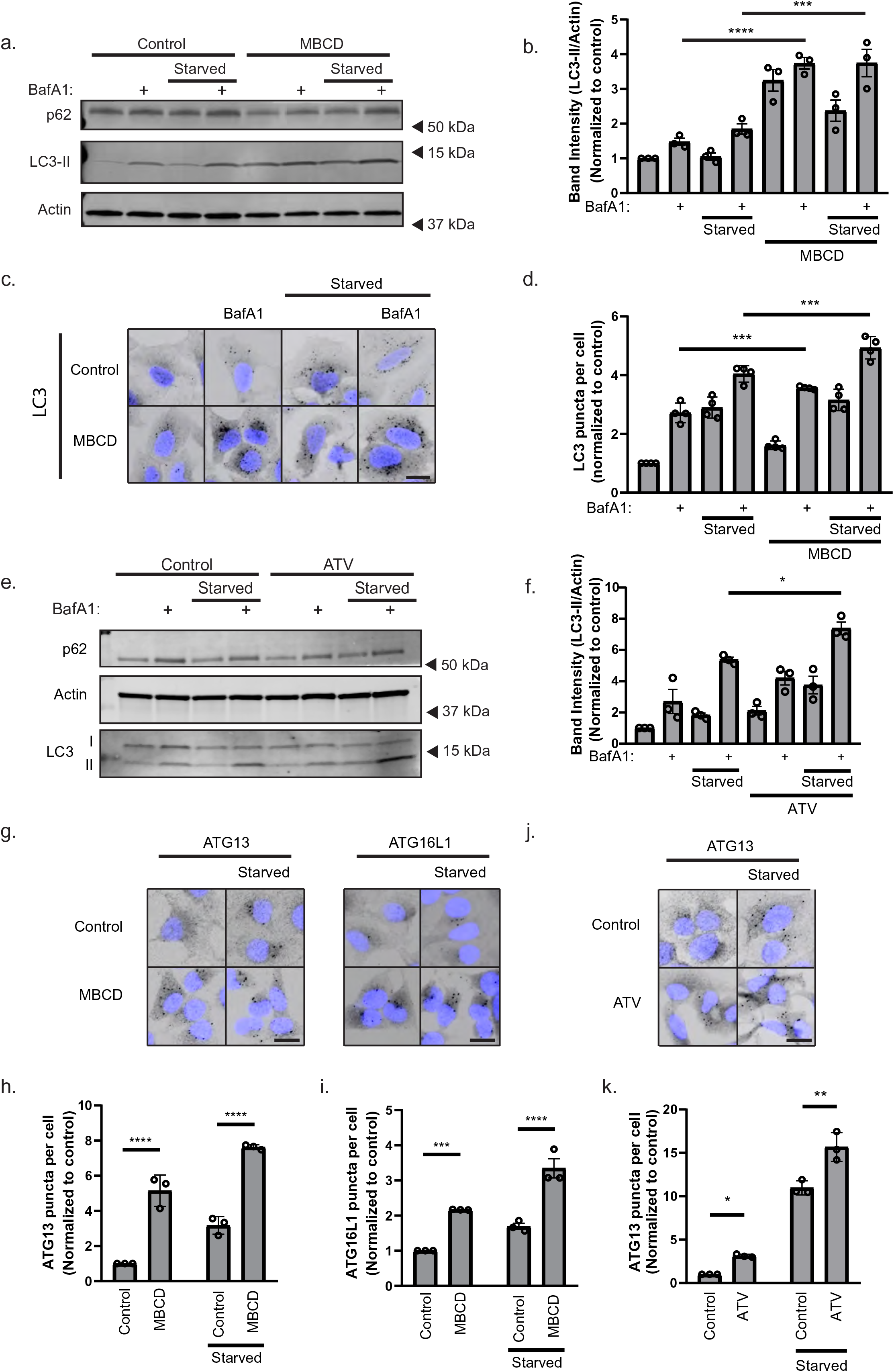
- Cholesterol depletion promotes autophagy initiation. **a** U2OS cells were treated with 2.5 mM MBCD in serum free DMEM or in EBSS supplemented ± 100 nM Bafilomycin A1 (BafA1) for 1 hr prior to western blot analysis with the indicated antibodies. **b** Quantification of LC3B-II band intensity relative to actin and normalized to DMEM control. Error bars = SEM. Significance was determined using 1-way ANOVA followed by Tukey’s comparisons test from n = 3 experiments. **c** U2OS cells were treated with 2.5 mM MBCD in serum free DMEM or in EBSS supplemented ± 100 nM BafA1 for 1 hr, prior to immunostaining with anti-LC3B antibody and widefield microscopy. Scale bar = 20µm. **d** Quantification of LC3B puncta number per cell normalized to the DMEM control. Error bars = SEM. Significance was determined using 1-way ANOVA followed by Tukey’s comparisons test from n = 4 experiments, >500 cells per condition. **e** U2OS cells were treated with 10 µM atorvastatin for 48 hrs, before amino acid starvation in EBSS 1 hr ± 100 nM BafA1, prior to western blot analysis with the indicated antibodies. **f** Quantification of LC3B-II band intensity relative to actin and normalized to DMEM control. Error bars = SEM. Significance was determined using 1-way ANOVA followed by Tukey’s comparisons test from n = 3 experiments. **g** U2OS cells were treated with 2.5 mM MBCD in serum free DMEM or EBSS for 1 hr and then immunostained with antibodies against ATG13 or ATG16L1. Scale bar = 20µm. **h-i** Quantification of data in g. ATG13 (h) and ATG16L1 (i) puncta number per cell were normalized to the DMEM control. Error bars = SEM. Significance was determined using 2-way ANOVA followed by Sidak’s comparisons test from n = 3 experiments, >500 cells per condition. **j** U2OS cells were treated with 10 µM atorvastatin for 48 hrs, before amino acid starvation in EBSS 1 hr and immunostaining with antibody against ATG13. Scale bar = 20µm. **k** Quantification of data in j. ATG13 puncta number per cell were normalized to the DMEM control. Error bars = SEM. Significance was determined using 2-way ANOVA followed by Sidak’s comparisons test from n = 3 experiments, >500 cells per condition. **** = p<0.0001, *** = p<0.001, ** = p < 0.01 and N.S. = not significant.

Autophagosome biogenesis is regulated by the autophagy initiation machinery comprising the ULK1 complex (ATG13, ULK1, FIP200, ATG101), PI3CK3 complex (ATG14L, BECN1, PIK3C3, P150) and the ATG12-ATG5-ATG16L1 complex. Given that LC3B lipidation was further increased in cells co-treated with MBCD and BafA1 compared to MBCD treated cells (Figure 1a-f), we suspected that the increase in LC3B lipidation was caused by changes in autophagosome biogenesis. In line with this, we observed a significant increase in the number of puncta positive for the early autophagy markers ATG13 and ATG16L1 in U2OS cells treated with MBCD for 1 hr, which was further increased upon amino acid starvation (Figure 1g-i). In support of this, ATV treated cells also exhibited enhanced ATG13 recruitment at both basal and starved states (Figure 1j-k). Taken together, these results show that cholesterol depletion promotes autophagy induction and enhances starvation-induced autophagy.

### Cholesterol depletion facilitates starvation-induced autophagy in an mTORC1 independent manner

Recent reports indicate that mTORC1 signaling is regulated by lysosomal cholesterol levels, and that lysosomal cholesterol depletion leads to the inactivation of mTORC1^29, 30^. Since we observed a synergistic effect on autophagy flux of cholesterol and amino acid starvation, we suspected that cholesterol depletion induces autophagy, in part, through a pathway that is independent of mTORC1 inactivation. To study this, we investigated the short-term temporal dynamics of mTORC1 signaling and autophagic flux in U2OS cells treated or not with either MBCD or ATV and starved at different time points (15, 30, 45 or 60 min) prior to immunoblotting for LC3B and the mTORC1 substrate p70S6K. Surprisingly, starvation-induced LC3B flux was significantly increased at 30 min in starved cells treated with MBCD, while this required 45 min in starved control cells (Figure 2a-b), indicating a more rapid induction of autophagosome biogenesis in cholesterol-depleted cells. This observation was recapitulated in ATV treated cells (Figure 2c-d). Interestingly, while the mTORC1 specific phosphorylation of p70S6K was completely lost after 15 mins of amino acid starvation, it took more than 30 mins in cells treated with MBCD (Figure 2e), indicating a delayed kinetic of mTORC1 inactivation in cells subjected to cholesterol depletion compared to that seen in starved cells. mTORC1 inactivation results in activation of the ULK1 complex^31^. Intriguingly, we found that ATG13 positive structures formed already after 15 min of MBCD treatment (Figure 2f-g), before the complete inactivation of mTORC1 (Figure 2e) suggesting that cholesterol depletion induces autophagy independent of mTORC1 signaling.

**Figure 2.**
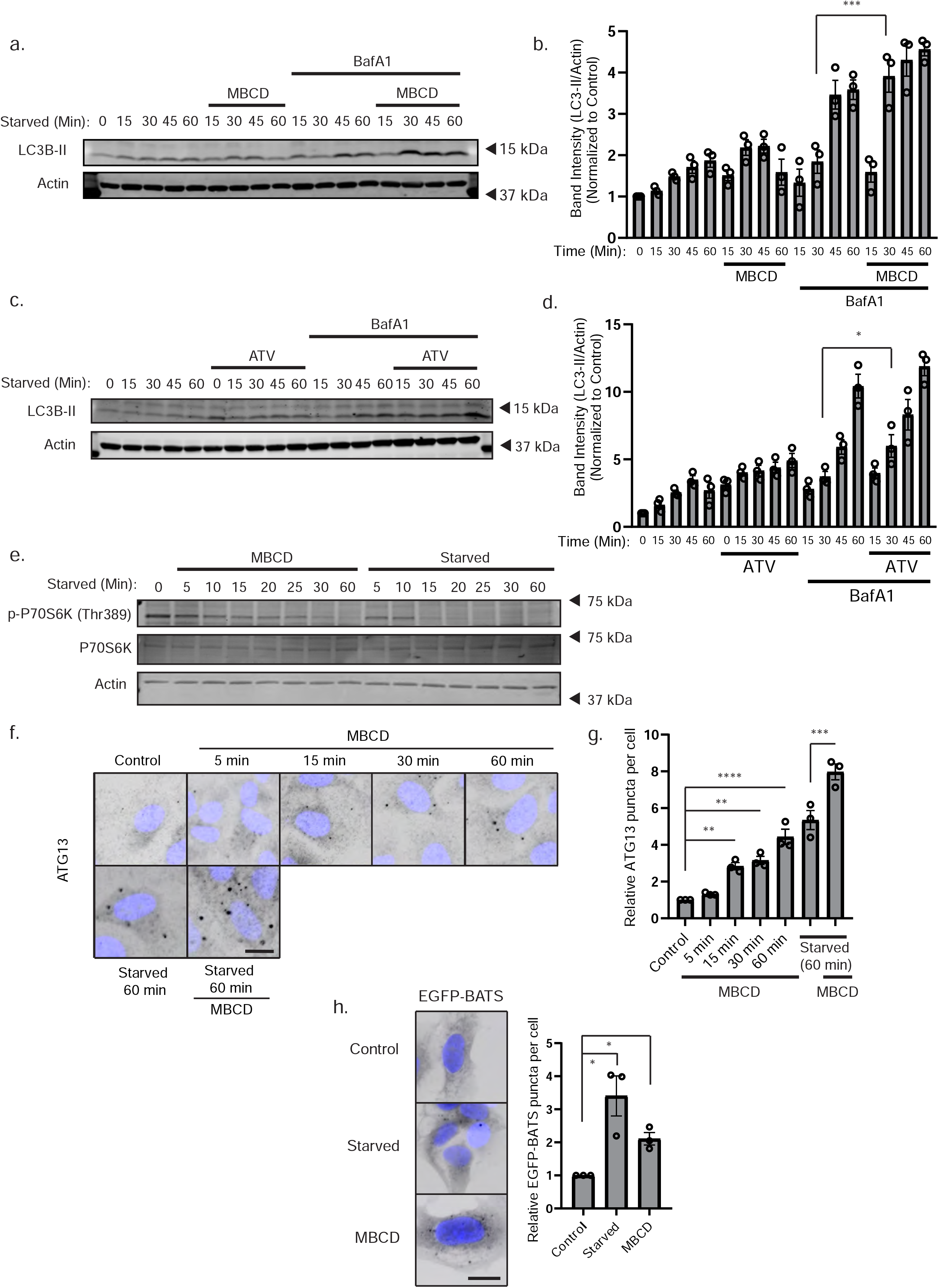
Cholesterol depletion alters starvation-induced autophagy dynamics in a mTORC1 independent manner. **a** U2OS cells were starved in EBSS and treated or not with 2.5 mM MBCD in DMEM ± 100 nM BafA1 for the indicated times before being lysed and subjected to western blot analysis. **b** Quantification of LC3B-II band intensity relative to Actin and normalized to DMEM control. Error bars = SEM. Significance was determined using 2-way ANOVA followed by Sidak’s comparisons test from n = 3 experiments. **c** U2OS cells were treated or not with 10 µM ATV for 48 hrs before starvation in EBSS ± BafA1 for the indicated times **d** Quantification of data in c. as LC3B-II band intensity relative to Actin and normalized to DMEM control. Error bars = SEM. Significance was determined using 2-way ANOVA followed by Sidak’s comparisons test from n = 3 experiments. **e** U2OS cells were treated for the indicated times with either 2.5 mM MBCD in DMEM or EBSS without MBCD before western blot analysis with the indicated antibodies. **f** U2OS cells were treated with 2.5 mM MBCD in DMEM or EBSS for the indicated time before fixation, immunostaining for ATG13 and widefield microscopy. Scale bar = 20 µm. **g** Quantification of ATG13 puncta per cell normalized to the control. Error bars = SEM. Significance was determined using 1-way ANOVA followed by Dunnett’s comparisons test from n = 3 experiments, >500 cells per condition. **h** Cells expressing EGFP-BATS were incubated in EBSS or serum free DMEM supplemented or not with 2.5 mM MBCD for 1 hr before fixation and widefield microscopy. Error bar = SEM. Scale bar = 20 µm. The number of EGFP-BATS puncta per cell was normalized to the DMEM control. Significance was determined using 1-way ANOVA followed by Dunnett’s from n = 3 experiments, >500 cells per experiment. **** = p<0.0001, *** = p<0.001, ** = p < 0.01 and N.S. = not significant.

As cholesterol is often found between the carbon chains of membrane phospholipids, we hypothesized that cholesterol removal promotes generation of curved membranes during *de novo* synthesis of autophagic membranes. To study this, we generated a membrane curvature reporter based on the amphipathic helix of the BATS domain of ATG14L1, known to bind to curved PtdIns(3)P-enriched membranes destined for autophagosome formation^32^ (Supplementary figure 1c). U2OS cells stably expressing EGFP-BATS formed puncta when starved for 1 hr (Figure 2h), where most puncta co-localized with autophagy initiation proteins (ATG13, ATG16L1, and WIPI2), as well as LC3B positive structures (Supplementary figure 1d), indicating that it marks early autophagy membranes. As predicted, MBCD treatment caused a significant increase of EGFP-BATS puncta compared to control cells (Figure 2h), indicating that cholesterol depletion induces the formation of curved early autophagy membranes.

### GRAMD1C is a negative regulator of starvation-induced autophagy

To elucidate whether genetic manipulation of cellular cholesterol levels also affects autophagy, we decided to deplete cholesterol transport proteins that are mediators of non-vesicular inter- organellar cholesterol movement at membrane contact sites^20^. The GRAM family of proteins, GRAMD1A, GRAMD1B, GRAMD1C, GRAMD2 and GRAMD3 (encoded by *GRAMD1A*, *GRAMD1B*, *GRAMD1C*, *GRAMD2a* and *GRAMD2b*) are ER anchored transmembrane proteins^22^ that function as cholesterol transport proteins, known to mediate plasma membrane to ER cholesterol transport^21^. Given that the GRAMs form a complex through their C-terminal region^21^ and since GRAMD1A has been implicated in autophagy^24^, we asked whether the other members of the GRAM family also have a role in autophagy. To study this, U2OS cells with stable inducible expression of the autophagy reporter mCherry-EGFP-LC3B (yellow-autophagosome, red- autolysosome due to quenching of EGFP in the acidic lysosome) were transfected with siRNA to individually deplete each GRAM family member, followed by quantification of the number of red- only puncta in starved and non-starved cells in the absence or presence of BafA1. Interestingly, there was a significant increase in the number of red-only puncta in GRAMD1C depleted cells compared to control, indicating that GRAMD1C is a negative regulator of starvation-induced autophagic flux (Figure 3a-b). Addition of BafA1 abolished the formation of red-only structures in both control and starved condition, confirming that these structures represent autolysosomes. In contrast to previous reports, GRAMD1A depletion did not inhibit starvation-induced autophagy, but rather promoted basal autophagy flux. The role for GRAMD1C in regulation of starvation-induced autophagy was validated by LC3B immunoblotting in cells depleted of GRAMD1C with two different siRNA oligos (Supplementary figure 2a). Indeed, the starvation- induced turnover of LC3B-II was significantly elevated in GRAMD1C knockdown cells, supporting a role for GRAMD1C as a negative regulator of autophagy (Figure 3c-d). We did not observe a significant change of p62 turnover upon GRAMD1C depletion (Supplementary figure 2b). Since lipidated LC3B and p62 do not fully represent the cargo diversity of the autophagosome and as LC3B is also implicated in non-canonical autophagy, we investigated the turnover of long-lived proteins, which are predominantly degraded through autophagy^33^ (Figure 3e). Indeed, the turnover of radioactively labelled long-lived proteins was increased in GRAMD1C depleted cells (Figure 3f), further demonstrating a role for GRAMD1C as a negative regulator of starvation- induced autophagy.

**Figure 3.**
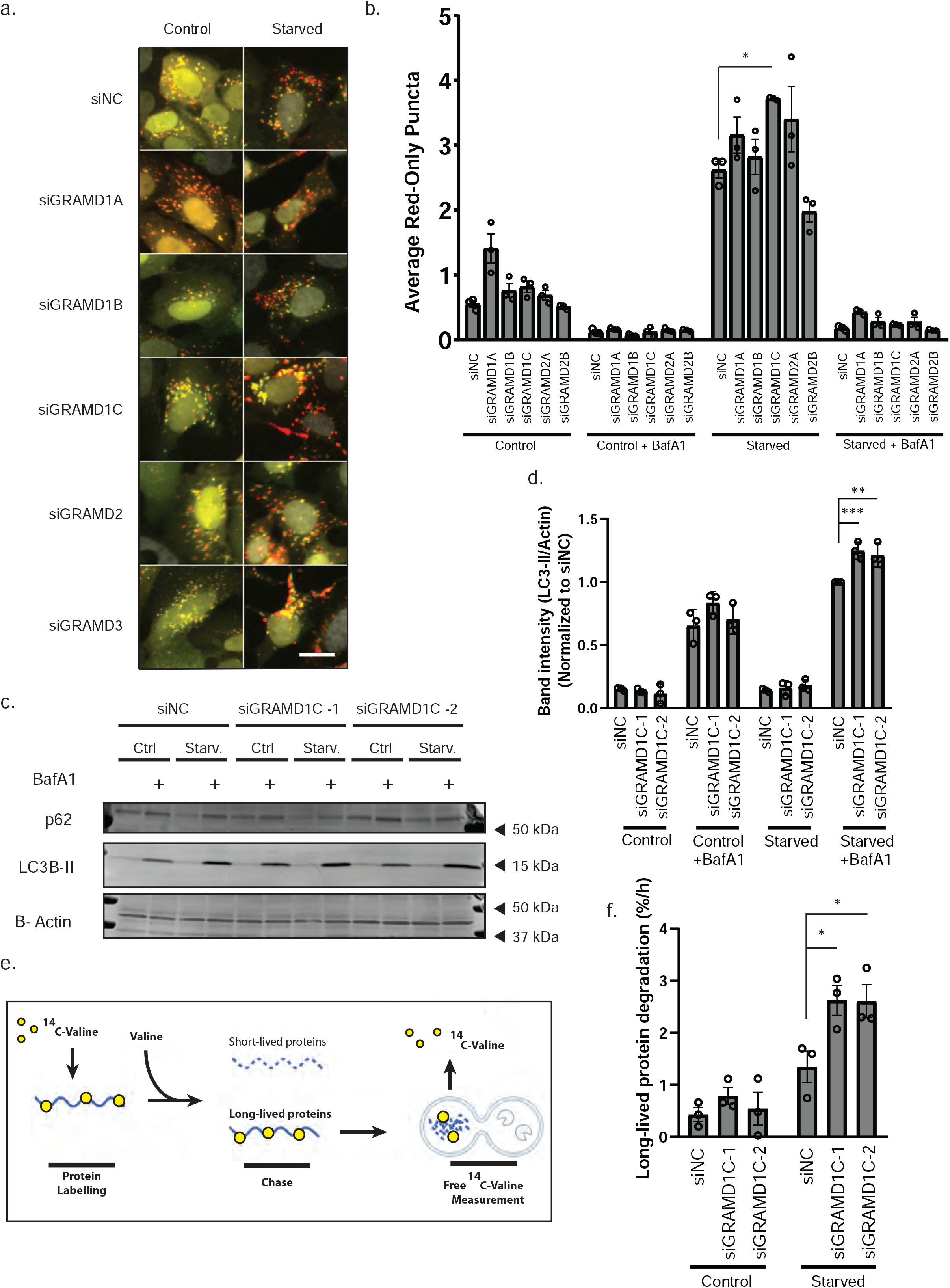
GRAMD1C is a negative regulator of starvation-induced autophagy. **a** U2OS cells stably expressing mCherry-EGFP-LC3B were transfected with siRNA against the indicated genes for 72 hrs before serum and amino acid starvation in EBSS for 2 hrs ± 100 nM BafA1 prior to fixation and widefield microscopy. Scale bar = 20 µm. **b** Quantification of data in a. The bars represent the average number of red-only puncta per cell. Significance was determined using 2-way ANOVA followed by Dunnett’s comparison test from n = 3 experiments, >500 cells per condition. Error bar = SEM. **c** U2OS cells were transfected with two different siRNA targeting GRAMD1C or a control for 72 hrs, followed by 2 hrs incubation in DMEM (Ctrl) or EBSS (Starv.) for 2 hrs ± 100 nM BafA1. The cells were then lysed and subjected to western blot analysis against the indicated proteins. **d** Quantification of band intensity of LC3B-II relative to Actin in c. Values are normalized to siNC starved + BafA1. Error bar = SEM. Significance was determined using 2-way ANOVA followed by Tukey’s comparison test from n = 3 experiments. **e** LLPD assay: Cells are incubated in culture media supplemented with ^14^C valine for 24 hrs, then washed and re-incubated in media containing non-radioactive valine for 16 hrs to allow degradation of short-lived proteins. The cells are then starved ± 100 nM BafA1 for 4 hrs, followed by analysis of radioactive ^14^C valine in the media and cells using a liquid scintillation counter. **f** U2OS cells were treated with the indicated siRNA and subjected to the LLPD assay. Significance was determined using 2-way followed by Tukey’s comparison test from n = 3 samples. *** = p<0.001, ** = p < 0.01 and N.S. = not significant.

To investigate whether GRAMD1C or any of the other GRAM family members regulate selective autophagy, U2OS cells with stable inducible expression of a mitophagy reporter construct (Mitochondria localization signal (MLS)-mCherry-EGFP) and stable ectopic Parkin expression were transfected with siRNA targeting each GRAM family protein, followed by induction of mitophagy by deferiprone (DFP) or CCCP. DFP is an iron chelator that induces a HIF1α-dependent response, leading to induction of Parkin-independent mitophagy^34^, while CCCP treatment results in loss of mitochondrial membrane potential and induction of Parkin-dependent mitophagy^35^. Cells were subjected to high content microscopy and quantification of red-only puncta, as a read- out for mitophagy. Knockdown of GRAMD1C neither affected Parkin-dependent (Supplementary figure 3a-b) nor Parkin-independent mitophagy (Supplementary figure 3c-d). Interestingly, GRAMD1A depletion increased DFP-induced mitophagy (Supplementary figure 3c-d). Taken together, our data suggest that GRAMD1C is a negative regulator of starvation-induced autophagy that is dispensable for selective clearance of mitochondria.

### GRAMD1C regulates autophagy initiation

As we found cholesterol depletion to increase autophagy initiation, we investigated if depletion of GRAMD1C caused a similar phenotype. Indeed, GRAMD1C depletion in U2OS cells resulted in increased membrane recruitment of several early autophagy machinery components, such as the ULK1 complex subunit ATG13, the ATG16L1-ATG5-ATG12 complex and the PtdIns(3)P effector protein WIPI2b^36^, as analyzed by quantification of the respective puncta from control and starved cells (Figure 4a-d). To corroborate these findings, we generated GRAMD1C knockout (GKO) cells (Supplementary figure 2c), and found a significant increase in ATG13 and ATG16L1 puncta formation in the GKO cells compared to passage matched wild type (Wt) cells, both before and after amino acid starvation (Figure 4e-g). Given that cholesterol depletion promoted the formation of BATS domain structures, we depleted GRAMD1C in cells expressing EGFP-BATS. Notably, GRAMD1C depletion promoted the recruitment of EGFP-BATS in starved cells (Figure 4h-i). Interestingly, we found GRAMD1C itself to also be delivered to the lysosomes upon starvation, as red-only puncta were seen in starved cells expressing mCherry-EGFP-GRAMD1C, which were sensitive to BafA1 (Supplementary figure 2d). Taken together, these results indicate that GRAMD1C regulates autophagosome biogenesis by removal of cholesterol from ER membranes, leading to recruitment of early autophagic markers.

**Figure 4.**
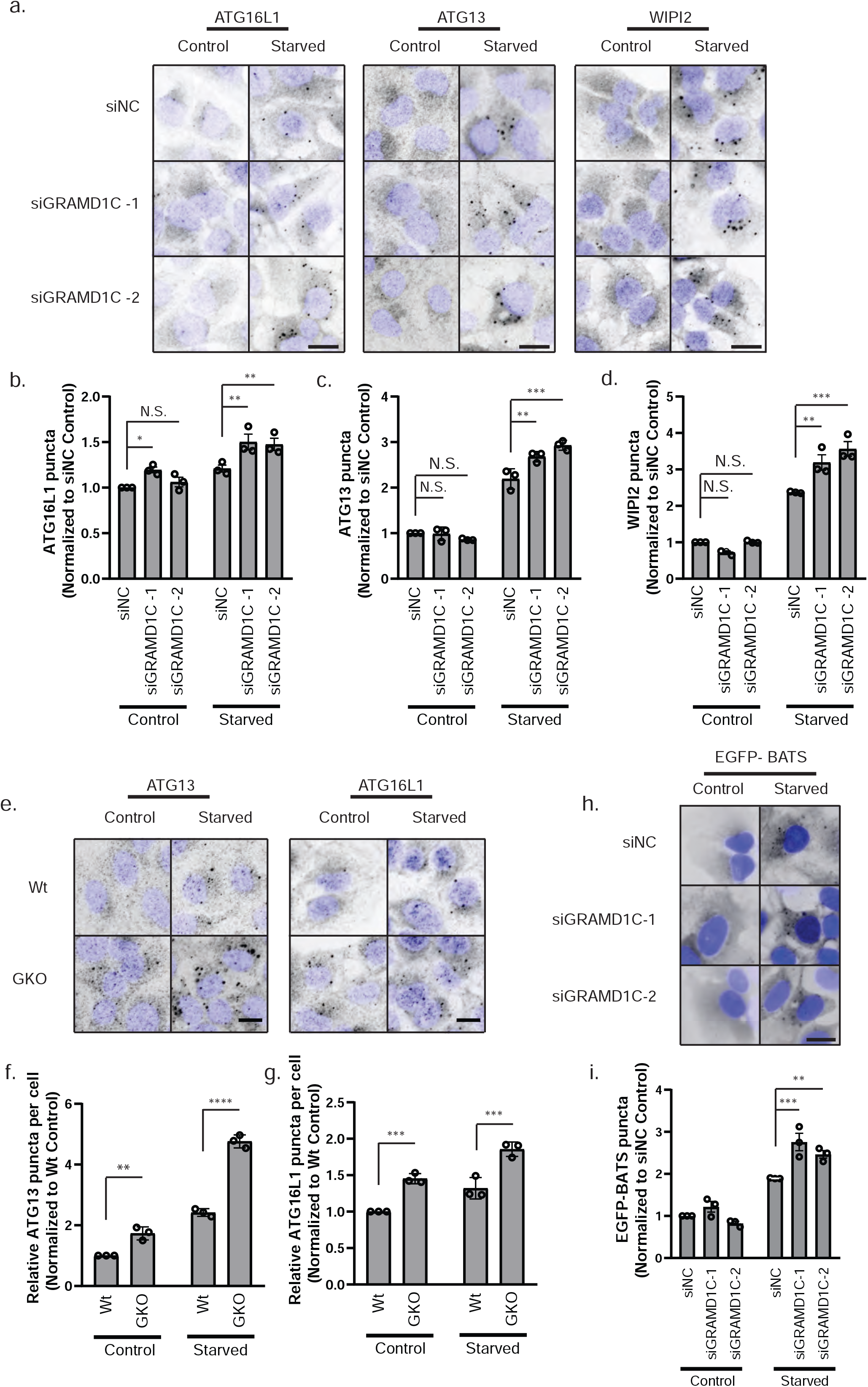
GRAMD1C regulates autophagy initiation during amino acid starvation. **a** U2OS cells were transfected with siRNA against control or GRAMD1C for 72 hrs prior to starvation in EBSS for 1 hr or incubation in DMEM (control). The cells were then fixed, immunostained with antibodies against ATG16L1, ATG13 or WIPI2 and subjected to widefield microscopy. Scale bar = 20µm. The number of **b** ATG16L1, **c** ATG13 and **d** WIPI2 puncta per cell were quantified and normalized to siNC control from n = 3 experiments, >500 cells per condition. Significance was determined using 2-way ANOVA followed by Tukey’s comparison test. Error bar = SEM. **e** Wild type (Wt) or GRAMD1C knockout (GKO) U2OS cells were starved or not in EBSS for 1 hr before immunostaining for ATG16L1 or ATG13. Scale bar = 20 µm. **f-g** The number of ATG13 (**f**) and ATG16L1 (**g**) puncta per cell were quantified and normalized to Wt control from n = 3 experiments, >500 cells per condition. Significance was determined using 2-way ANOVA followed by Sidak’s comparison test. **h** U2OS cells expressing EGFP-BATS were transfected with siRNA against control or GRAMD1C for 72 hrs before starvation in EBSS for 1 hr. Scale bar = 20 µm **i** The number of EGFP-BATS puncta per cell was quantified and normalized to siNC control from n = 3 experiments, >500 cells per condition. Significance was determined using 2-way ANOVA followed by Tukey’s comparison test. Error bar = SEM. **** = p<0.0001, *** = p<0.001, ** = p < 0.01 and N.S. = not significant.

### The GRAM domain of GRAMD1C mediates its interaction to the mitochondria

The yeast orthologue of the GRAMs, Lam6, is promiscuously enriched at different organellar contact sites, such as the vCLAMP (vacuole and mitochondria patch), NVJ (nuclear vacuolar junction) and ERMES (ER-mitochondria encounter structures)^37^. In contrast, GRAMD1C has only been reported to be recruited to the plasma membrane to facilitate cholesterol import^22, 38^. The lipid-binding GRAM domain is thought to be responsible for protein targeting to a specific organelle and given the relative conservation of the GRAM domain of GRAMD1C to that of Lam6 (37.88% sequence identity) (Supplementary figure 4a), we asked whether GRAMD1C also exhibited other contact site localizations. Stable cell lines expressing EGFP-tagged wild type GRAMD1C or the GRAM domain only (Figure 5a) were analyzed by live cell microscopy. As expected, GRAMD1C-EGFP localized to the ER, but was also found to be enriched at regions of ER/mitochondria overlap (Figure 5b). Interestingly, the EGFP-GRAM domain localized to structures that appeared to associate onto mitochondria for a few seconds before dissociating, suggesting that the GRAM domain of GRAMD1C facilitates a transient interaction with mitochondria (Figure 5c). Indeed, the mitochondrial interaction of the GRAM domain was validated by isolation and immunoblotting of mitochondria from cells expressing the EGFP-GRAM domain, showing the EGFP-GRAM domain both in the crude and pure mitochondria fractions (Figure 5d). In an attempt to identify proteins interacting specifically with the GRAM domain of GRAMD1C, and by extension characterize the interactome of GRAMD1C, we carried out co-immunopurification coupled mass-spectrometry (coIP-MS) analysis of the interactome of EGFP tagged GRAMD1C and GRAMD1C lacking the GRAM domain (ΔGRAM). As expected, GRAMD1A, GRAMD1B, GRAMD2 and GRAMD3 were amongst the interactome of GRAMD1C (Figure 5e, Table I). Strikingly, GO-term enrichment revealed that GRAMD1C mainly interacted with proteins of mitochondrial origin (Figure 5f, Table I). Additionally, several mitochondrial proteins such as NDUFAF2, SHDB and ATAD3A, as well as ER-mitochondria contact site proteins, VDAC1 and ACSL4 were enriched in the interactome of GRAMD1C, but absent in that of ΔGRAM (Figure 5g), indicating that the mitochondrial interaction of GRAMD1C is dependent on the GRAM domain. As the interaction between mitochondria and ER can be affected by changes in mitochondrial structure, we analyzed mitochondrial morphology upon the depletion of the GRAMs, but did not see any changes to mitochondrial morphology upon depletion of GRAMD1C (Supplementary figure 4b-c). Thus, our results show that GRAMD1C interacts with the mitochondria through its GRAM domain.

**Figure 5.**
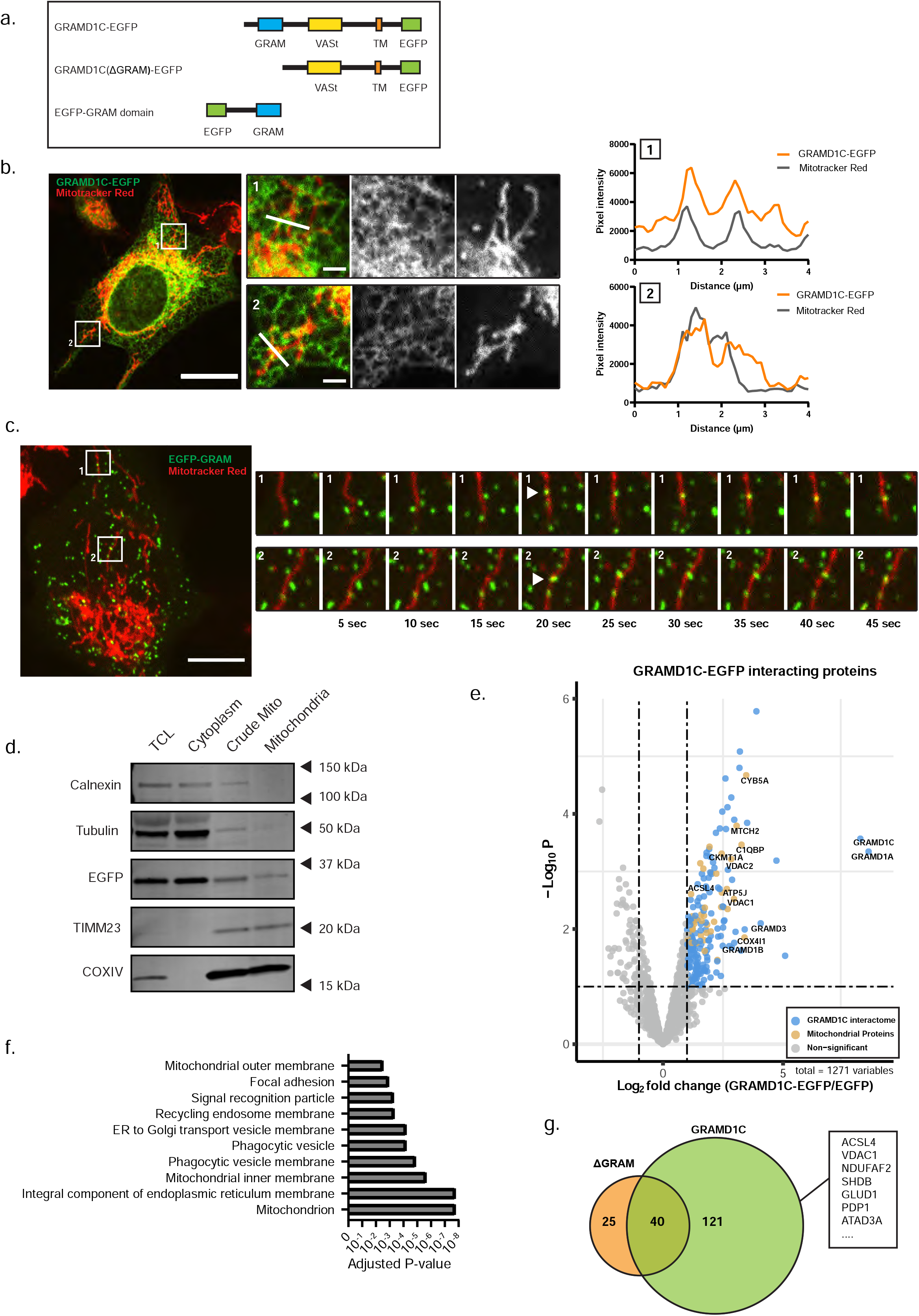
GRAMD1C interacts with the mitochondria through the GRAM domain. **a** Overview of the EGFP-tagged GRAMD1C constructs used. **b** U2OS cells stably expressing GRAMD1C-EGFP were stained with Mitotracker Red and subjected to live cell confocal microscopy. The graph depicts the pixel intensity along the white line drawn in the inset. Scale Bar = 20 µm, inset = 2 µm. **c** U2OS cells stably expressing EGFP-GRAM domain were stained with Mitotracker Red and subjected to live cell confocal microscopy. The insets represent snapshots showing recruitment of EGFP-GRAM to mitochondria. Scale Bar = 10 µm. **d** Mitochondria were isolated using percoll density centrifugation from U2OS cells expressing EGFP-GRAM and subjected to western blot analysis for the indicated proteins. TLC: total cell lysate. **e** GRAMD1C- EGFP was immunopurified and co-purified proteins were identified using mass spectrometry analysis. The interactome of GRAMD1C-EGFP was compared to the interactome of EGFP. Significant hits (p<0.05) are colored blue or brown (mitochondrial). **f** GO cellular compartment enrichment of proteins co-purified with GRAMD1C-EGFP. **g** Venn diagram of interacting proteins of GRAMD1C-EGFP and GRAMD1C(ΔGRAM)-EGFP. The proteins listed are examples of mitochondrial and mitochondria-ER contact site proteins.

### GRAMD1C regulates mitochondrial bioenergetics

Given the ability of GRAMD1C to interact with mitochondria, we next asked if GRAMD1C can potentially regulate cholesterol movement between the ER and the mitochondria. For that reason, we developed a method for mitochondrial cholesterol quantification based on the addition of recombinant mCherry-tagged cholesterol binding domain of Perfringolysin O (mCherry-D4)^39^ to isolated mitochondria. As expected, MBCD treatment decreased mCherry-D4 binding to purified mitochondria, indicating that mCherry-D4 selectively binds to cholesterol on isolated mitochondria (Figure 6a-b). Importantly, increased levels of mCherry-D4 were detected on the mitochondria isolated form of GRAMD1C KO cells compared to control cells (Figure 6a-b), indicating that GRAMD1C regulates mitochondrial cholesterol levels. In support of this, cholesterol oxidase-based quantification of mitochondrial cholesterol revealed a similar increase of mitochondrial cholesterol in GRAMD1C KO cells compared to control cells (Figure 6c). At the same time, loss of GRAMD1C caused a reduction of ER cholesterol levels, as seen through an increase in SREBP target gene expression (Supplementary figure 4d), in line with a previous observations^40^. Additionally, the abundance of cholesterol-associated proteins (STARD9, ERLIN, SQLE, NPC2, and APOB) in GRAMD1C depleted cells were altered as seen through proteomic analysis of siGRAMD1C treated cells (Supplementary figure 4e, Table II). Thus, our data indicate that GRAMD1C facilitates cholesterol transport from mitochondria to the ER.

**Figure 6.**
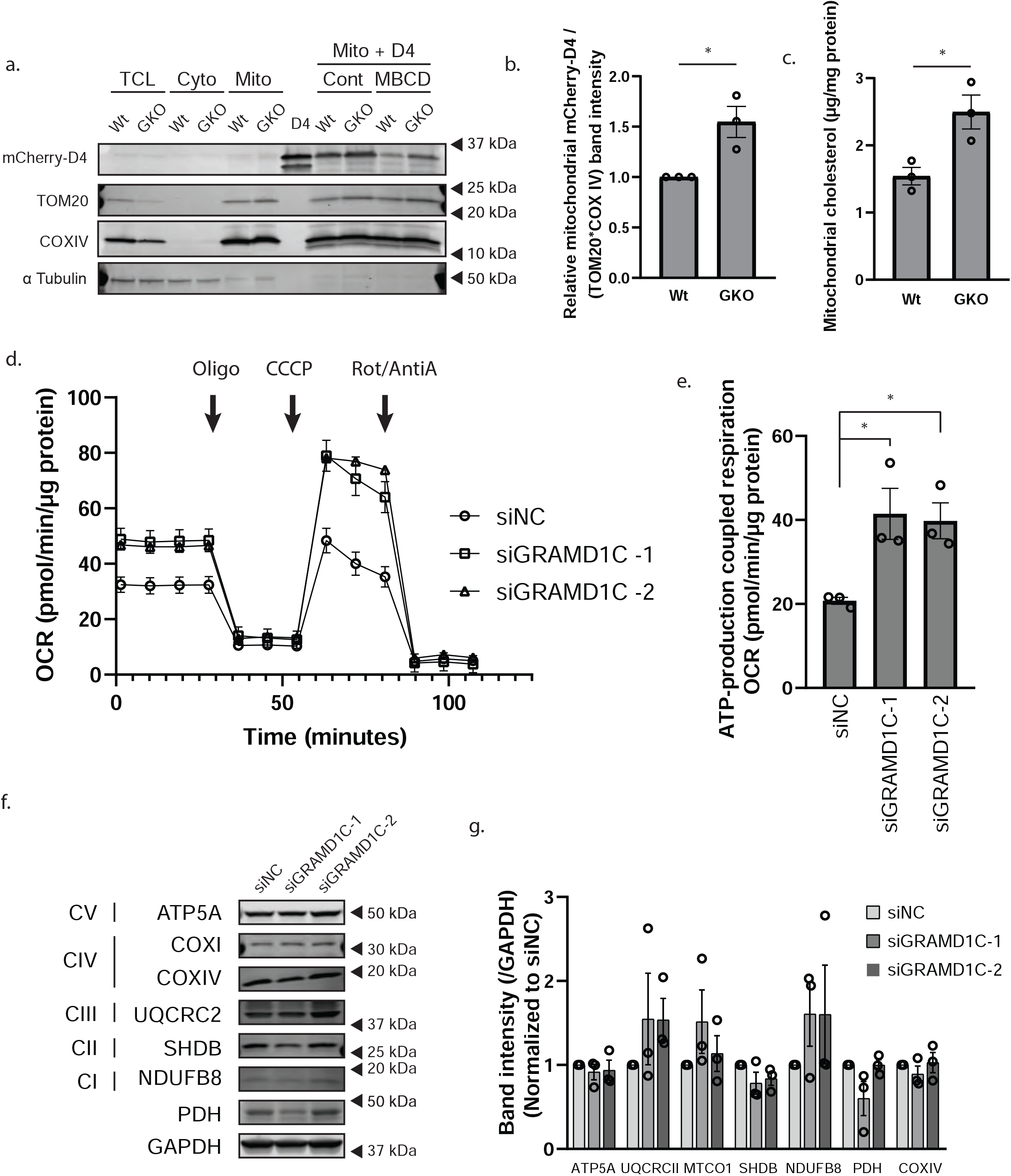
GRAMD1C regulates mitochondrial bioenergetics. **a** Wt and GRAMD1C knockout (GKO) cells expressing 3XHA-EGFP-OMP25 were treated or not with MBCD, followed by isolation of mitochondria using affinity purification of the HA-tagged OMP25 from a total cell lysate (TCL). The isolated mitochondria were incubated with the cholesterol probe mCherry-D4 (recombinant protein shown in lane 7), before washing and western blot analysis with the indicated antibodies. **b** Quantification of mCherry-D4 band intensity in the isolated mitochondria fractions from a. relative to the average band intensity of the mitochondrial proteins TOM20 and COXIV and normalized to wt cells. Significance was determined using Students T-test from n =3 experiments. Error bar = SEM. **c** Mitochondria from Wt or GKO cells were isolated and mitochondrial lipids were extracted. The lipids were then oxidized with cholesterol oxidase, which generates H2O2 that reacts with a colorimetric probe. Absorbance changes corresponding to cholesterol abundance were normalized to mitochondrial protein concentration. Significance was determined using Student’s T-test from n = 3 experiments. Error bar = SEM. **d** Mitochondrial oxygen consumption was analyzed in control and GRAMD1C knocked down cells using the Seahorse analyzer. Oxygen consumption was measured after gradual addition of Oligomycin, CCCP and Rotenone/Antimycin A. **e** ATP-linked respiration is calculated from the difference between the maximal respiratory capacity and the proton leak. Significance was determined using 1-way ANOVA followed by Bonferroni’s comparison test from n = 3 experiments. Error bar = SEM. **f** Cell lysates from control (siNC) and siGRAMD1C treated cells were subjected to western blot analysis with antibodies against the indicated OXPHOS components. **g** The graph represents the quantification of band intensities of data in f. relative to GAPDH and normalized to siNC. Significance was determined with 1-way ANOVA followed by Tukey’s comparison test from n =3 experiments. Error bar = SEM. **** = p<0.0001, *** = p<0.001, ** = p < 0.01 and N.S. = not significant.

In order to investigate the implication of this increased mitochondrial cholesterol level, we inspected mitochondrial function in GRAMD1C depleted cells. Interestingly, GRAMD1C depletion increased the ATP-production linked respiration and the maximal respiratory capacity as analyzed by Seahorse XF Analyzer (Figure 6d-e). Furthermore, western blot analysis did not reveal significant changes to the OXPHOS proteins in GRAMD1C knockdown cells (Figure 6f-g), suggesting that the change in mitochondrial respiration was not caused by changes in the mitochondrial proteome. Similarly, mitochondrial membrane potential and total cellular reactive oxygen species (ROS) were not altered in GRAMD1C knockdown cells (Supplementary figure 4f- g). In summary, our results indicate that GRAMD1C is a negative regulator of mitochondrial cholesterol abundance and mitochondrial bioenergetics.

### GRAM family expression is prognostic in ccRCC

GRAM proteins have previously been implicated in tumorigenesis, as GRAMD1B depletion was found to promote breast cancer cell migration^41^, while *GRAMD1C* transcript levels seems to positively correlate with the level of immune cell infiltration and overall survival in Clear Cell Renal Carcinoma (ccRCC) patients^42^. ccRCC is a type of kidney cancer that stems from the epithelial cells of the proximal convoluted tubule of the kidney^43^, and is characterized by altered mitochondrial metabolism and aberrant lipid and cholesterol accumulation^44^. Given that *GRAMD1C* expression correlates with overall survival in ccRCC^42^ and forms a heteromeric complex with the other GRAMS^21^, we investigated the involvement of the complete GRAM family in ccRCC using tumor gene expression data from the TCGA KIRC cohort^45^. Interestingly, the expression of several GRAM family members was significantly associated with survival outcome. Similar to *GRAMD1C*, high *GRAMD2B* expression was associated with improved patient survival in ccRCC. By contrast, low expression of GRAMD1A and GRAMD1B was favorable with respect to survival (Figure 7a). We further found a weak negative correlation between *GRAMD1C* expression and *GRAMD1A* and *GRAMD1B* levels (Supplementary figure 5a), reflecting their differential influence on overall survival. Furthermore, while *GRAMD1C* expression was decreased in advanced stage tumors, the expression of *GRAMD1A* and *GRAMD1B* showed an opposite pattern, with increased expression in late stage tumor samples compared to early stage tumor samples (Supplementary figure 5b).

**Figure 7.**
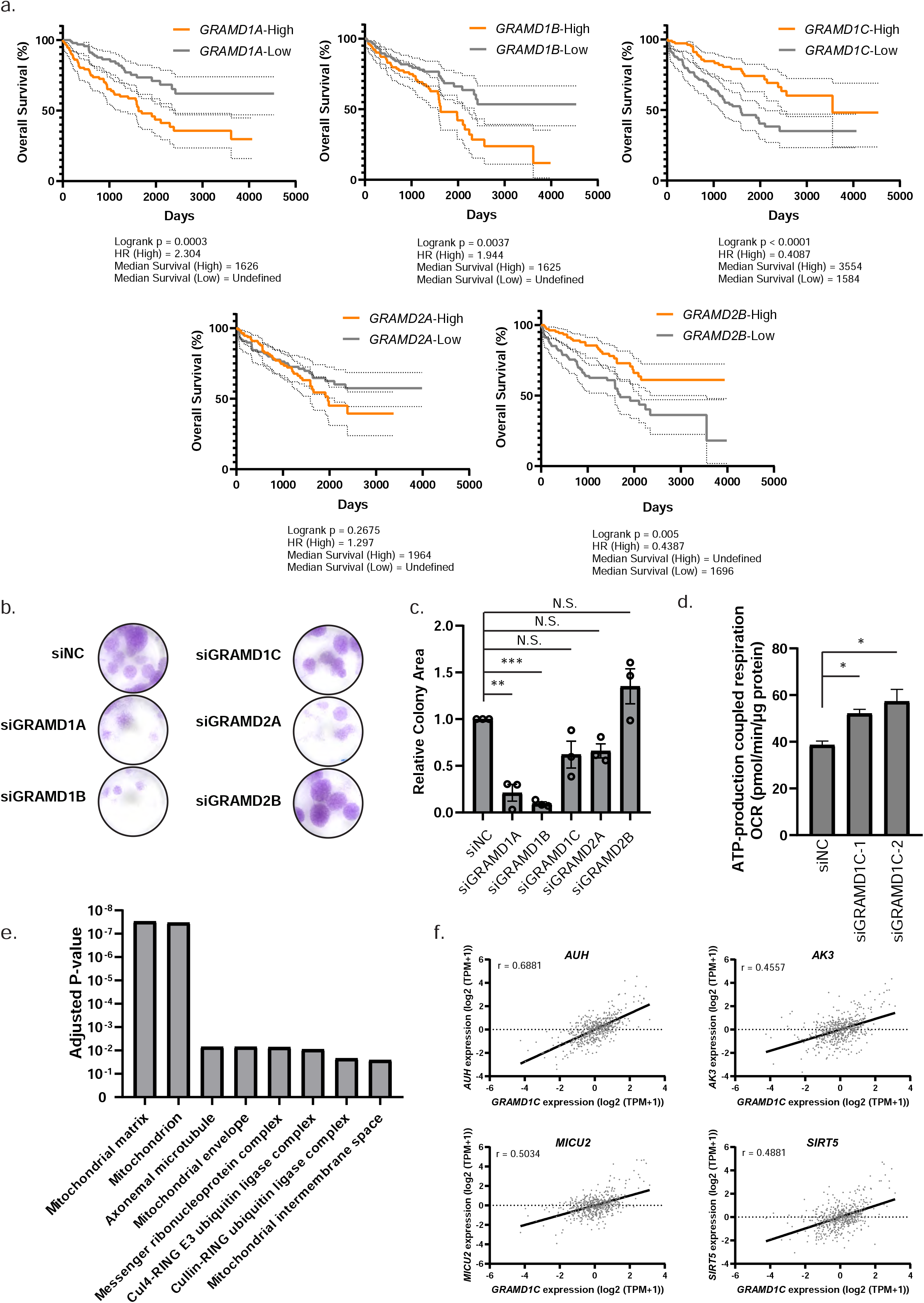
The GRAMs are involved in ccRCC survival. **a** Samples from the KIRC TCGA study were stratified based on *GRAM* expression. Overall survival of samples with high *GRAM* expression (Upper quartile, orange line) were compared to low *GRAM* expression (Lower quartile, gray line). The dotted lines represent 95% confidence interval. P-values were obtained using Log-rank (Mantel-Cox) test. **b** For the colony formation assay, 786- O cells were treated with the indicated siRNAs and incubated for 3 weeks prior to fixation and staining with crystal violet stain. **c** Colony area was quantified and normalized to siNC. Significance was determined using 1-way ANOVA followed by Dunett’s comparison test from n = 3 experiments. **d** ATP-linked respiration, calculated from the difference between the maximal respiratory capacity and the proton leak from Seahorse analysis, in 786-O cells treated with control (siNC) or siRNA against GRAMD1C for 72 hrs. Significance was determined using 1-way ANOVA followed by Dunnett’s comparison test. Error bar = SEM. **e** Top 100 co-expressed genes of *GRAMD1C* in the KIRC TCGA dataset were downloaded from GEPIA2.cancer-pku.cn, and subjected to GO Cellular Compartment enrichment using Enrichr^83, 84^. **f** Top four co-expressed mitochondrial genes with *GRAMD1C* in the KIRC TCGA cohort plotted against the expression of GRAMD1C. *** = p<0.001, ** = p < 0.01 and N.S. = not significant.

We were able to validate this observation using a colony formation assay in 786-O ccRCC cells transfected with siRNA against all GRAMs, showing that depletion of *GRAMD1A* and *GRAMD1B* significantly decreased the ability of 786-O cells to form colonies (Figure 7b-c), supporting the role of these genes in ccRCC survival. As overall survival is also affected by invasive and migration capabilities of the tumor, we analyzed migration of 786-O cells depleted of GRAMD1A-C using a wound healing assay. However, only the migration of siGRAMD1B treated cells was slightly decreased (Supplementary figure 6a-b). Similar to our observation in U2OS cells, *GRAMD1C* depletion promoted ATP-production linked respiration in ccRCC 786-O cells (Figure 7d), indicating that the relationship between mitochondrial bioenergetics and GRAMD1C is conserved among cell lines.

In order to better understand the role of GRAMD1C in ccRCC, we analyzed the genes co-expressed with *GRAMD1C*, as co-expressed gene networks can allow identification of functionally related genes^46^. Interestingly, *GRAMD1C* is co-expressed with several mitochondrial genes in ccRCC samples, including AUH, AK3, MICU2 and SIRT5, which all moderately correlated with GRAMD1C with Pearson’s correlations values of above 0.45 (Figure 7e-f). In conclusion, members of the GRAM family contribute to the regulation of overall survival of ccRCC patients, possibly through modulation of metabolism and cancer cell survival.

## Discussion

In this study, we investigated the effects of cholesterol and the cholesterol transport protein, GRAMD1C on starvation-induced autophagy. We show that cholesterol depletion promotes autophagy initiation and enhances starvation-induced autophagy flux independently of mTOR signaling. Additionally, we show that depletion of GRAMD1C promotes starvation-induced autophagy. Importantly, we find GRAMD1C to interact with mitochondria through its PH-like GRAM domain but has no effect on Parkin-dependent or -independent mitophagy. Depletion of GRAMD1C leads to increased mitochondrial cholesterol abundance and increased mitochondrial bioenergetics. Finally, we identify the GRAM family as genes involved in ccRCC survival, highlighting the pathophysiological relevance of cholesterol transport proteins.

Autophagosomes are small vesicles (approx. 0.5-1 um) formed *de novo* from ER-associated sites ^47^. In order to facilitate the generation of the autophagosome, forming membranes will require a high degree of curvature and flexibility, which must be partially supported by specialized lipid compositions. Reflecting this, localized fatty acid synthesis has been shown to occur at autophagy biogenesis sites and are required for autophagosome generation^48–50^. Furthermore, the autophagy proteins ATG2 and ATG9 have been shown to facilitate lipid delivery to growing autophagosomes^51–53^. The role of cholesterol during autophagosome biogenesis remains to be clarified, but given that membrane cholesterol increases membrane rigidity, it is likely that high cholesterol abundance at autophagosome initiation sites is unfavorable for autophagosome biogenesis.

Interestingly, cholesterol depletion has a synergistic effect on autophagic flux in response to amino acid starvation, suggesting that cholesterol and amino acid depletion activate autophagy in part through mutually exclusive mechanisms. Indeed, our data suggest that the short-term effect of cholesterol depletion on autophagy induction is independent of mTORC1 inactivation and rather is mediated by a change in the membrane curvature, as membrane recruitment of the curvature-sensing BATS domain of ATG14L was significantly increased in cells depleted of cholesterol. As ATG14L is recruited to PtdIns(3)P enriched ER-associated omegasomes upon induction of autophagy by starvation^54^, we speculated that a removal of cholesterol at such sites might facilitate autophagosome biogenesis.

The proteins of the GRAM family are ER anchored transmembrane proteins that function as cholesterol transport proteins^21^ that previously have been shown to be recruited to the PM in order to facilitate ER cholesterol import^22^. We demonstrate here that GRAMD1C depletion promotes starvation-induced autophagy by increasing the numbers of autophagosome initiation sites. Most interestingly, mitochondrial cholesterol levels were increased in GRAMD1C depleted cells, suggesting that it regulates cholesterol transport between mitochondria and the ER. Supplementing these findings, we show that GRAMD1C interacts with the mitochondria through its GRAM domain and co-precipitates with mitochondrial proteins. Interestingly, instead of its GRAM domain, the interaction of GRAMD1B to the mitochondria was reported to be dependent on a mitochondrion targeting sequence located upstream of its GRAM domain^55^, indicating that the GRAMs may be differently recruited to target organelles. It remains to be shown whether GRAMD1C interacts directly with any mitochondrial proteins or whether mitochondria fragments were indirectly pulled down with GRAMD1C. Interestingly, GRAMD1C copurified with ACSL4 and VDAC1, both markers of ER-mitochondria contact sites, supporting the localization of GRAMD1C to these regions. We were not able to establish if GRAMD1C is a *bona fide* contact tether ^56^, or if this interaction is transient in nature. Autophagy initiation occurs at ER-mitochondria contact sites^57, 58^, thus placing GRAMD1C at sites of autophagosome biogenesis and we propose a model where GRAMD1C functions as a negative regulator of autophagy by regulating cholesterol levels at autophagosome initiations sites (Supplementary figure 6c).

Recent reports suggest the involvement of GRAMD1A in the regulation of autophagosome biogenesis. Surprisingly, GRAMD1A depletion in U2OS cells did not inhibit autophagy as previously reported in MCF7 and HEK293T cells^24^, but rather enhanced basal autophagy flux. While it was recently suggested that GRAMD1C negatively regulates autophagy in immortalized mouse myoblast C2C12 cells, this statement was not supported with autophagy flux experiments^59^. Nevertheless, these observed differences could stem from the divergence in the cholesterol requirement between cell types. The effects of GRAMD1C depletion on autophagy were not as drastic as compared to MBCD-mediated cholesterol depletion, leading us to suspect that GRAMD1A and GRAMD1B were able to partially compensate for the loss of GRAMD1C given their highly similar structures. However, GRAMD1C KO (GKO) cells exhibited a more pronounced induction of autophagy initiation events, suggesting that GRAMD1C is not completely redundant in the long-term. Furthermore, GRAMD1C did neither alter Parkin-dependent nor -independent mitophagy, possibly reflecting a difference in the *de novo* formation of the autophagosome during selective and non-selective autophagy.

The increased mitochondrial cholesterol levels seen in GRAMD1C depleted cells is reminiscent of the increase in mitochondrial cholesterol in Niemann Pick C1 (NPC1) depleted cells^60^. However, the relationship between mitochondrial cholesterol and respiration is not clear. Previous studies have found that mice fed with a cholesterol-enriched diet displayed increased mitochondrial cholesterol and decreased mitochondrial respiration ^61^, but cholesterol removal with MBCD^62^ and simvastatin also decreased mitochondrial respiration^63^. This discrepancy can possibly be attributed to the different experimental models and cholesterol loading/depletion systems used. Our results suggest that mitochondrial cholesterol accumulation caused by the loss of GRAMD1C promotes oxidative phosphorylation. While we have not been able to establish a mechanism for this, proteomic analysis of GRAMD1C depleted cells suggest an altered composition of proteins involved in cellular metabolism. Moreover, the abundance of Glycogen Synthase Kinase 3 beta (GSK3B), Glycogenin-1 (GYG1) and Glycerol-3-phosphate phosphatase (PGP), proteins involved in glycogen synthesis, were decreased in GRAMD1C depleted cells. In contrast the glycolysis regulator 6-phosphofructo-2-kinase/Fructose-2,6-bisphosphatase 4 (PFKFB4) was increased in cells treated with siGRAMD1C (Supplementary figure 4e). The significance of these changes is not clear, but it suggests that metabolic rewiring accompanies the loss of GRAMD1C.

ccRCC cells exhibit a disrupted cholesterol homeostasis, accumulating up to 8 times more higher cholesterol compared to normal kidney cells^64, 65^. However this increase does not appear to stem from increased cholesterol synthesis and possibly originates from aberrant cholesterol transport and metabolism^64^. It is therefore interesting that high expression of GRAMD1C and GRAMD2B was found to associate with improved survival of ccRCC patients, while the opposite was found for GRAMD1A and GRAMD1B. This suggests that the GRAMs may have opposing roles in ccRCC carcinogenesis and survival despite their domain similarities. Critically, mirroring the observation on overall survival, we found that depletion of *GRAMD1A* and *GRAMD1B* in 786-O ccRCC cells caused a significant decrease in cell survival, suggesting that the GRAMS are relevant therapeutic targets for ccRCC.

In conclusion, our results show that short-term cholesterol depletion is favorable for autophagosome biogenesis by increasing the membrane recruitment of early core autophagy proteins. We show that depletion of the ER-anchored cholesterol transport protein GRAMD1C promotes starvation-induced autophagy and find GRAMD1C to interact with mitochondria to facilitate mitochondria-ER cholesterol transport. Finally, we find that the expression of various *GRAM* genes correlates with ccRCC survival. These results underline the importance of cholesterol transport proteins in autophagy and mitochondrial bioenergetics and warrants further investigation into the regulation of membrane cholesterol during autophagosome biogenesis and cancer.

## Materials and Methods

### Antibodies

The following primary antibodies were used: anti-LC3B (Western blotting, #3868, Cell Signaling), anti-LC3B (Immuno-fluorescence microscopy, #PM036, MBL), anti-p62 (#610833, BD biosciences), anti-tubulin (#T5168, Sigma), anti-actin (#3700, Cell Signaling), anti-EGFP (#632381, Takara), anti-mCherry (#PA534974, Thermo Fisher), anti-TOM20 (#17764, Santa Cruz), anti- TIM23 (#611223, BD Biosciences), anti-COX IV (#4850, Cell Signaling), anti-ATG13 (#13468, Cell Signaling), anti-ATG16L1 (#PM040, MBL), anti-WIPI2 (#Ab105459, Abcam) anti-VDAC1 (#ab15895, Abcam total OXPHOS antibody (#ab110413, Abcam), anti-PDH (#2784S, Cell Signaling), GAPDH (#5174, Cell Signaling), p70S6K (#9202, Cell Signaling), phospho-P70S6K (Thr389) (#9205, Cell Signaling).

Secondary antibodies for western blotting used were anti-mouse DyLight 680 (#SA5-10170, Thermo Fisher), anti-rabbit DyLight 800 (#SA5-10044, Thermo Fisher). Secondary antibodies used for immunofluorescence were Anti-rabbit Alexa Fluor 488 (#A-21206, Invitrogen) and Anti-mouse CY3 (#115-165-146, Jackson).

### Materials

The following chemicals were used: Bafilomycin A1 (#BML-CM110, Enzo Life Sciences), CCCP (#BML-CM124, Enzo Life Sciences) Oligomycin A (#S1478, Selleckchem). Antimycin A (#A8674, Sigma Aldrich), DFP (#37940, Sigma Aldrich), DTT (#441496p, VWR), Rotenone (#R8875, Sigma Aldrich), MBCD (#M7439, Sigma Aldrich). For lysis buffers, Complete EDTA-free protease inhibitor (#05056489001, Roche) and PhosStop phosphatase inhibitor (#04906837001, Roche) were used. For live cell imaging, Mitotracker Red (#M22425, Thermo Fisher) was used. For measurement of ROS, CellRox (#C10422, Thermo Fisher) was used. To measure mitochondrial membrane potential, cells were incubated in TMRE (#T669, Thermo Fisher). For amino acid starvation, cells were cultured in Earle’s Balanced Salt Solution (EBSS) (Invitrogen). Percoll (#sc-500790A, Santa Cruz). For high-throughput widefield microscopy, cells were cultured in µ-Plate 96 Well ibiTreat (#89626, Ibidi).

Silencer Select siRNA (Thermo Fisher) were used against the following target genes. GRAMD1A (s33529), GRAMD1B (s33113), GRAMD1C (s29400 siGRAMD1C-1, s29401 siGRAMD1C-2), GRAMD2a (s47069), GRAMD2b (s35302), OPA1 (s9851), DRP1 (s19559) and Negative Control (s813).

### Cell lines

U2OS, U2OS Flp-In and HEK293T were cultured in complete DMEM containing 10 % v/v fetal bovine serum and 100 U/mL Penicillin + 100 µg/mL Streptomycin at 37°C in 5% CO2. 786-O cells were grown in RPMI supplemented with containing 10 % v/v fetal bovine serum and 100 U/mL Penicillin + 100 µg/mL Streptomycin at 37°C in 5% CO2. U2OS TRex FlpIn cells (kindly provided by Steve Blacklow, Harvard Medical School, US) were used for generation of stable inducible cell lines.

### Lentivirus production and stable cell line generation

Stable cells were generated using lentiviral transduction and Flp-In Trex system^66^. Target genes were cloned into pLenti-III or pLVX viral expression vectors which were co-transfected with psPAX2 and pCMV-VSVG into HEK293FT cells to generate lentiviral particles. The lentiviral particles were then concentrated using Lenti-X. The resulting lentivirus solution was added to cells and supplemented with 8µg/mL polybrene. The cells were then selected using the appropriate antibiotics (Puromycin (#p7255, Sigma Aldrich) or Zeocin (#R25001, Invitrogen)).

### Knockout cell line generation and validation

GRAMD1C knockout cells were generated using the PX459 system^67^. In short, U2OS cells were transfected with the PX459 vector expressing guides against GRAMD1C. Guide sequences were designed using CHOPCHOP^68^. 24 hrs post transfection, transfected cells were selected with 3 µg/ml puromycin for 72 hrs. Single cell clones are then selected using limited dilution into 96 well tissue culture plates. Due to a lack of an antibody that recognizes endogenous GRAMD1C, validation of the knockout clones was done by sequencing of the relevant region of GRAMD1C from genomic DNA. A total of 15 sequencing reactions were done, all of which indicated a frameshift mutation (E245S*fs20) (Supplementary figure 2b).

### siRNA knockdown

siRNA mediated knockdown was preformed using reverse transfection of siRNA against the target gene at a final concentration of 10nM per oligonucleotide. siRNAs were delivered using Lipofectamine RNAi max (Invitrogen). After 24 hrs, the cells were washed and replenished with normal media. Cells stably expressing inducible mCherry-EGFP-LC3b and MLS-mCherry-EGFP were supplemented with media containing 100 ng/ml Doxycycline. At 72 hrs, the cells are then treated as described in the figure legends.

Due to the lack of reliable antibodies for endogenous GRAMD1C, knockdown was validated using qPCR against GRAMD1C (Supplementary figure 2a).

### ROS measurement

U2OS cells were transfected as described above. Upon 48 hrs post transfection, cells were plated in 12-well plates and left in the incubator O/N. Upon 72 hrs post transfection cells were treated with CellRox (#C10422, Thermo Fischer) for 10 minutes according to manufacturer’s instructions. After washing cells were trypsinised, washed in PBS twice and analyzed using the BD™ LSR II flow cytometer. A total of three experiments in duplicates were performed and fluorescent signal was analyzed using FlowJo.

### Wound healing assay

768-O renal carcinoma cells were transiently transfected as described above. Transfected cells were seeded at 96 well-plates (#4379, Essen Bioscience) 48 hrs post transfection. A total of 4x10^4^ cells/well were seeded in triplicates at approximately 100% well-density. At 72 hrs post transfection one scratch per well was made using the Incucyte® 96-well WoundMaker Tool (#4563, Sartorius). Plate was then loaded in the Incucyte incubator. One image every 20 minutes for a total of 24 hrs was acquired for each well. Results were analyzed using the Integrated Cell Migration analysis module (#9600-0012, Sartorius).

### Microscopy and sample preparation

Cells were seeded on glass coverslips or onto glass bottom 96 well imaging plates and treated as indicated. The cells were then washed twice in prewarmed PBS prior to the addition of warmed fixation solution (3.7% PFA, 200 mM HEPES pH 7.1) and incubated at 37°C for 20 mins. The fixed cells were then washed 3 times with PBS. Cells destined for immunofluorescence staining were permeabilized with 0.2% NP-40 in PBS for 5 mins. The cells were then washed twice in PBS and incubated in 5% BSA in PBS for 30 mins. The cells were incubated at 20°C for 1h in primary antibody diluted in 5% BSA in PBS. The cells were then washed 3 times with PBS and incubated in secondary antibody diluted in 5% BSA in PBS at 20°C for 45 mins. The samples were then washed with PBS. Coverslips were mounted on cover slides using Prolong Diamond Antifade Mounting Solution and wells of the 96-well imaging plate were filled with PBS to prevent cells from drying out.

Quantitative spot counting of ATG13, ATG13, WIPI2 and LC3 immunostained cells was carried out using a Zeiss AxioObserver microscope (Zen Blue 2.3 Zeiss) fitted with a 20x Objective (NA 0.5). The samples were illuminated using a solid-state light source (Colibri 7) and multi-bandpass filters (BP425/30, 534/50, 688/145). Imaging of cells expressing MLS-mCherry-EGFP, and mCherry-EGFP-LC3b, was done using the ImageXpress Micro Confocal (Molecular Devices) using a 20x objective (NA 0.45). Confocal images were take using the Zeiss LSM 800 microscope (Zen Black 2012 SP5 FP3, Zeiss) equipped with at 63x oil immersion objective (NA 1.4). Samples were illuminated using a laser diode (405nm), AR-Laser Multiline (458/488/514nm), DPSS (561nm) and HeNe-Laser (633nm). Live cell confocal imaging was done with cells in a humidified chamber at 37°C supplemented with 5%CO2 on the Dragonfly (Oxford Instrumentals) with a 60x objective (NA 1.4) using a EMCCD camera. For live cell imaging, the cells were treated as indicated in the figure legend, before replacing the culture media with FluoroBrite DMEM (#A1896701, Thermo Fisher).

### Bioimage analysis

ATG16, ATG13, WIPI2 and LC3 puncta were quantified using the CellProfiler software (2.2.0, 3.1.9 and 4.07, Broad Institute)^69, 70^. The nuclei were determined using manual thresholding and object identification of the nuclear stain, and the cells were defined based on a set distance from the center of the nuclei and was confirmed by comparing to the background cytosolic staining of the other channels. Puncta were determined using manual thresholding, object enhancement and object identification. For analysis of mCherry-EGFP-LC3b and mCherry-EGFP-MLS cells, red-only structures were determined by weighting the red signal to match the green signal and by dividing the weighted red signal by the green signal using the CellProfiler software (2.2.0, 3.1.9 and 4.07, Broad Institute). Values that are larger than 1 will represent mitochondria/LC3 structures that have a stronger red signal compared to the green signal. The resulting analysis was manually compared to the image to confirm the accuracy of the imaging pipeline. A value of 1.5 corresponds to twice the signal of red compared to the green.

### cDNA synthesis and RT-PCR

RNA was isolated using the RNeasy plus kit (Qiagen) according to manufacturer’s instructions. RNA integrity was confirmed by agarose gel prior to cDNA synthesis. cDNA was synthesized using SuperScript II reverse transcriptase (Thermo Fisher) and real time quantitative PCR was carried out using SYBR Green Real Time PCR master mix (Qiagen). Normalization of target genes were done against TATA-box-binding protein (TBP) using the 2^-ΔΔCt^ method.

### Western blotting

Cells were treated as indicated in the figure legends before being washed twice in ice cold PBS. The cells were then lysed in NP-40 lysis buffer (50 mM HEPES pH 7.4, 150 mM NaCl, 1 mM EDTA, 10 % glycerol, 0.5 % NP-40, Phosphatase inhibitor and Complete Protease inhibitor Cocktail (Roche)). The protein concentration of the lysates was measured with BCA assay (Thermo Fisher). The lysates were run on an SDS-PAGE at 20-30 µg of protein per well before transfer to a PVDF membrane. Blocking was done using a PBS blocking solution (Licor). The resulting membrane was then incubated using the specified primary and secondary antibodies. Visualization of the bound far-red secondary antibodies was performed using the Odyssey CLx imaging system (Licor), and densitometric quantification was performed using the ImageStudio Lite software (Licor).

### Oxygen consumption rate measurement

U2OS cells resuspended in complete DMEM were seeded into Seahorse XFe24 Cell Culture microplates at a concentration of 3.5 x 10^4^ cells per well. The plate was incubated in a humidified incubator at 37°C for 12 hr. The media was then replaced with DMEM without Sodium Bicarbonate (pH 7.4) before analysis with the Seahorse XFe24 Analyzer according to manufacturer’s instructions (XF mito stress test, Agilent). DMEM containing specific mitochondrial inhibitors were loaded into the injector ports of the Seahorse Sensor Plates to obtain the following final concentrations per well (CCCP: 1 µM, Oligomycin: 1.5 µM, Rotenone: 0.5 µM, Antimycin A: 0.5 µM). After the analysis, the cells were washed in ice cold PBS and lysed for protein quantification using BCA Assay (Thermo Fisher). Quantification was conducted on the Seahorse Analytics software (seahorseanalytics.agilent.com, Agilent), using the measured protein concentration from each well for normalization.

### Mitochondria isolation

Mitochondria were isolated using two different methods. For percoll density gradient isolation, cells are scraped with ice cold mitochondrial isolation buffer (5 mM Tris-HCl pH 7.4, 210 mM mannitol, 70 mM Sucrose, 1 mM EDTA, 1 mM DTT, 1X PhosStop, 1x PIC) and mechanically lysed using a cell homogenizer (Isobiotech) equipped with a 16 µm clearance ball by passing the cell suspension 10 times through the homogenizer. The resulting solution was then centrifuged at 1500 xg for 5 mins at 4 °C to pellet nucleus and unbroken cells. The suspension was then centrifuged at 14000 xg for 20 mins to obtain a crude mitochondrial pellet. The pellet was then resuspended in mitochondrial isolation buffer and layered above a premade percoll gradient of 50 %, 22 % and 15 % in a 5 ml ultracentrifuge tube. The tube was then centrifuged at 30000 xg for 1 h. A white layer between the 50 % and 22 % gradient is isolated using a syringe and needle. Percoll was separated from the isolated mitochondrial fraction by washing in mitochondrial isolation buffer and centrifugation at 14000 xg for 15 mins for 4-5 times. After the final wash, the pellet containing isolated mitochondria was lysed with RIPA lysis buffer.

For affinity purification of mitochondria, mitochondria were isolated from cells stably expressing 3xHA-EGFP-OMP25 according to Walter et al^71^ with minor modifications. In short, cells were scraped in ice cold KPBS (136 mM KCl, 10mM KH2PO4, pH 7.25) and mechanically lysed using a cell homogenizer (Isobiotech) equipped with a 16 µm clearance ball by passing the cell suspension 10 times through the homogenizer. The resulting solution is then centrifuged at 1500 xg for 5 mins at 4 degrees to pellet nucleus and unbroken cells. The supernatant was then incubated with anti-HA magnetic beads (Thermo Fisher) for 5 mins, before washing with KPBS and resuspension in 2x SDS Page loading buffer.

### Cholesterol quantification

Cholesterol quantification was done using the Cholesterol / Cholesteryl Ester Assay Kit (Abcam, ab65359). Briefly, cells were washed twice in ice cold PBS, scrapped, and spun down. The resulting cell pellet was then resuspended with Chloroform:Isopropanol:NP-40 (7:11:0.1) to extract lipids. The mixture was then air dried at 50 °C to remove the chloroform. The resulting lipids were analyzed according to manufacturer’s instructions. The resulting values were normalized to proteins measured by BCA assay.

### Mitochondrial cholesterol mCherry-D4 assay

Isolated mitochondria were incubated in Mitochondria Isolation Buffer 100 µg/mL of mCherry- D4 ± 5 mM MBCD at 37 degrees for 30 mins. The mitochondria were then washed 3 times in mitochondria isolation buffer and lysed with 2x SDS-page loading buffer and immediately subjected to western blot analysis.

### Mitochondrial structure classification

Cells stably expressing IMLS were treated with siRNA against the GRAMs, OPA1 and DRP1. After 72 hrs of knockdown, the cells were fixed and imaged. In these cells, only DAPI and the EGFP signals were measured. The mitochondrial intensity distribution, texture, shape, and area was measured using CellProfiler. The results were used in CellProfiler Analyst (v2.2.1, Broad Institute)^72, 73^ to classify mitochondrial morphology. Mitochondria of siOPA1 and siDRP1 cells represented fragmented and tubular phenotypes respectively. The classifier was trained with a confusion matrix >0.90 for each phenotype.

### Long-lived protein degradation

Cells were incubated in complete DMEM supplemented with 0.25 µCi/mL L-^14^C-valine (Perkin Elmer) for 24 hr. The radioactive media was then removed, and the cells were washed three times with complete DMEM supplemented with 10mM L-Valine, and finally chased for 16 hr in complete DMEM supplemented with 10 mM L-valine. The cells were then washed three times in PBS and either starved in EBSS or not for 4 hrs, in the presence or absence of 100 nM BafA1. The supernatant was collected into tubes containing 15% trichloroacetic acid before subsequent incubation at 4°C for 12 hr. The cells remaining in the dish were lysed with 0.2 M KOH. The supernatant was recovered by centrifugation. The supernatant and the cell lysate were added into separate scintillation tubes containing Ultima Gold LSC cocktail (Perkin Elmer) and the radioactivity was measured by a TriCarb 3100TR liquid scintillation counter (Perkin Elmer). Long-lived protein degradation was calculated by dividing the radioactivity in the supernatant fraction by the total radioactivity in both the supernatant and cell lysate.

### Co-IP Mass spec

Cells expressing GRAMD1C-EGFP or GRAMD1C(ΔGRAM)-EGFP were lysed with NP-40 lysis buffer (150 mM, 1.0% NP-40, 50 mM Tris-HCl pH 8.0) supplemented with PhosStop phosphatase inhibitor (Sigma) and complete protease inhibitor cocktail (Sigma). Co-immunopurification was then carried out using EGFP-TRAP (Chromotek) according to manufacturer’s instructions. The resulting beads with the coprecipitated proteins were washed twice with 50mM ammonium bicarbonate. Proteins on beads were reduced and alkylated and further digested by trypsin for overnight at 37 degree. Digested peptides were transferred to new tube, acidified and the peptides were de-salted for MS analysis.

### LC-MS/MS

Peptides samples were dissolved in 10ul 0.1% formic buffer and 3 ul loaded for MS analysis. LC-MS/MS analysis of the resulting peptides was performed using an Easy nLC1000 liquid chromatography system (Thermo Electron, Bremen, Germany) coupled to a QExactive HF Hybrid Quadrupole-Orbitrap mass spectrometer (Thermo Electron) with a nanoelectrospray ion source (EasySpray, Thermo Electron). The LC separation of peptides was performed using an EasySpray C18 analytical column (2 µm particle size, 100 Å, 75 μm inner diameter and 25 cm ; Thermo Fisher Scientific). Peptides were separated over a 90 min gradient from 2% to 30% (v/v) ACN in 0.1% (v/v) FA, after which the column was washed using 90% (v/v) ACN in 0.1% (v/v) FA for 20 min (flow rate 0.3 μL/min). All LC-MS/MS analyses were operated in data-dependent mode where the most intense peptides were automatically selected for fragmentation by high-energy collision-induced dissociation.

Raw files from LC-MS/MS analyses were submitted to MaxQuant 1.6.17.0 software^74^ for peptide/protein identification. Parameters were set as follow: Carbamidomethyl (C) was set as a fixed modification and PTY; protein N-acetylation and methionine oxidation as variable modifications. First search error window of 20 ppm and mains search error of 6 ppm. Trypsin without proline restriction enzyme option was used, with two allowed miscleavages. Minimal unique peptides were set to one, and FDR allowed was 0.01 (1%) for peptide and protein identification. The Uniprot human database was used. Generation of reversed sequences was selected to assign FDR rates. Further analysis was performed with Perseus^75^, limma^76^, Package R^77^. Volcano plots were plotted with EnhancedVolcano^78^.The gene ontology (GO) analysis was performed with shinyGO ^79^.

### TCGA data

Survival data and cancer stage from all samples in the TCGA-KIRC (clear cell renal cell carcinoma, data release 27.0) cohort was downloaded with the TCGABiolinks R package^80^. Survival curve comparisons were carried out in GraphPad Prism 8.0.1 using Log-rank (Mantel-Cox) test.

### Sequence alignment

The GRAM domain sequences of each GRAM protein were obtained from Uniprot, which were then aligned using Clustal Omega^81^ and BlastP^82^.

### Crystal violet staining

786-O cells were seeded into 6 well and 24 well plates in quadruplicates and treated as indicated. After 3 weeks, the cells are fixed in staining solution (6% Glutaraldehyde, 0.5% Crystal Violet) for 1 hr at room temperature. The fixation solution was removed, and the cells were rinsed by multiple gentle immersion in H2O. The stained cells were then imaged on a BioRad ChemiDoc MP analyzer. Quantification of colony area was done using ImageJ software.

### Statistics and significance

Statistical analysis was carried out using Prism (8.01) using the test as indicated in the figure legends. All relevant statistical tests are described in the figure legends and all data values come from distinct samples. **** = p>0.0001, *** = p >0.001, ** = p>0.01, * = p > 005 or N.S. = not significant.

### Primers used in this study

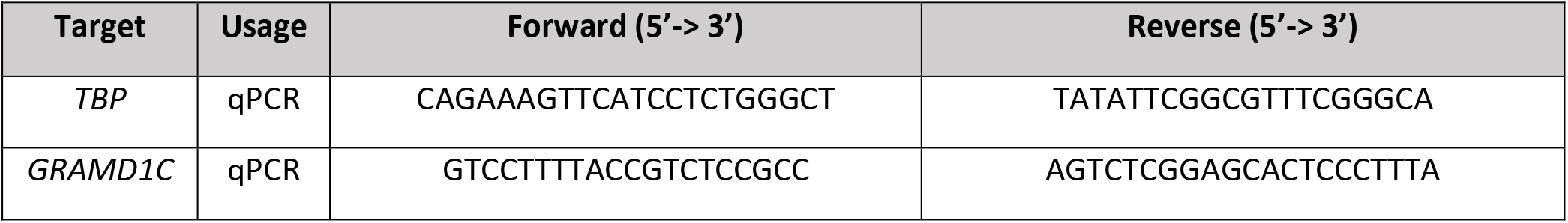
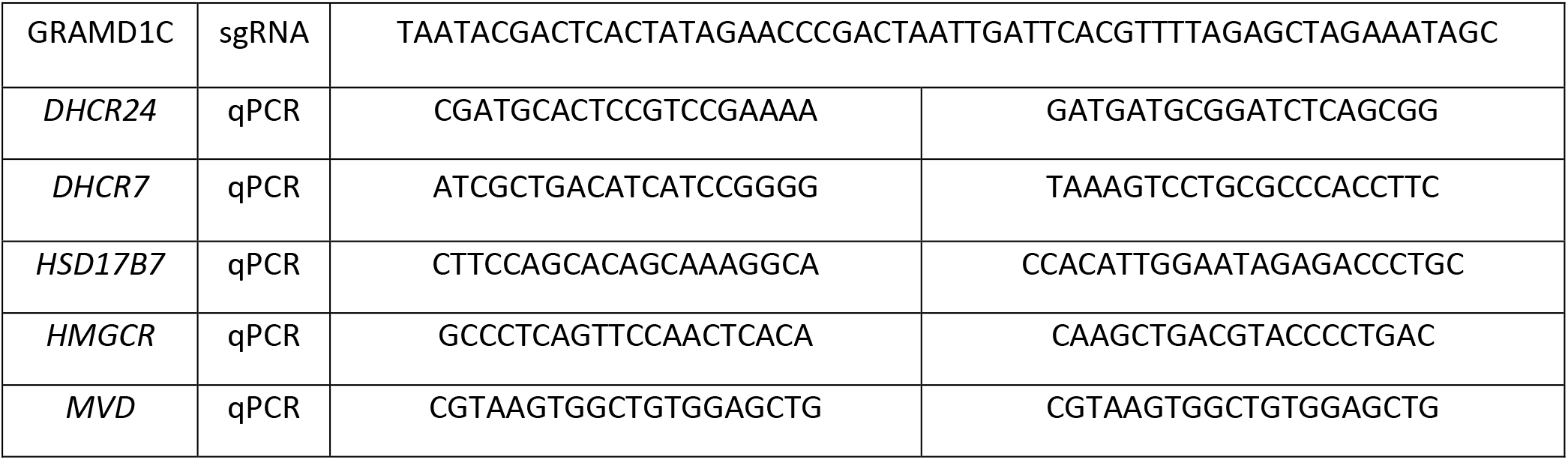

### Plasmids used in this study

| Vector | Description | Source |
| --- | --- | --- |
| pENTR3C | Entry vector | Invitrogen |
| pENTR3C-GRAMD1C | Cloning human GRAMD1C from cDNA from U2OS into a pENTR3C backbone. | This study |
| pENTR3C-GRAMD1C (ΔGRAM) | Cloning human GRAMD1C lacking the GRAM domain into a pENTR3C backbone. | This study |
| pLenti-PGK-GRAMD1C-EGFP | C-terminal EGFP tagged GRAMD1C expressing lentiviral vector generated using Gibson Assembly. | This study |
| pLenti-PGK-GRAMD1C (ΔGRAM)-EGFP | C-terminal EGFP tagged GRAMD1C (ΔGRAM) expressing lentiviral vector generated using Gibson Assembly | This study |
| pCMV-VSV-G | Envelope protein vector for lentivirus production. | Gift from Bob Weinberg (Addgene # 8454) |
| psPAX2 | Lentiviral packaging vector for lentivirus production. | Gift from Didier Trono (Addgene # 12260) |
| pLenti-PGK-EGFP-GRAM domain | EGFP-GRAM domain cloned from entry vector into a pLenti-PGK vector using Gibson Assembly. | This study |
| pcDNA5-MLS-EGFP-mCherry | Vector for inducible mitophagy reporter using the FlpIn system. | This study |
| pLenti-PGK-3xHA-EGFP-OMP25 | 3xHA-EGFP-OMP25 vector used for affinity purification of mitochondria. The insert was cloned from <sup>71</sup>. | This study |
| pLenti-PGK-EGFP-BATS | EGFP-BATS domain cloned from ATG14L plasmid into a pLenti-PGK vector using Gibson Assembly. | This study |
| pDest-FlpIn-mCherry-EGFP-GRAMD1C | Doxycycline inducible mCherry-EGFP-GRAMD1C construct generated using gateway cloning. | This study |

## Acknowledgements

This work was supported by the Research Council of Norway through its Centre of Excellence funding scheme (Project: 262652) and FRIPRO grant (Project: 221831), the Norwegian Cancer Society (Project: 171318), the Marie Skłodowska-Curie (MSC) ETN grant under the European Union’s Horizon 2020 Research and Innovation Programme (Grant Agreement No 765912 DRIVE) and the UiO Scientia Fellow program through the MSC scheme – Co-funding of Regional, National and International Programmes (COFUND). The authors would like to thank Sonia Peña Pérez for help with the graphical abstract and Laura Rodriguez de la Ballina for technical help and constructive feedback. The authors would also like to thank Sachin Singh and the Proteomics Core Facility for their assistance with the mass spectrometry based proteomic experiments and the Norwegian Core Facility for Human Pluripotent Stem Cells at the Norwegian Center for Stem Cell Research for mycoplasma testing of our cells.

## Author contributions

M.N., C.C., A.S conceived and planned the experiments. S.S. carried out the analysis of mass spec data. M.N., C.C., A.L. and L.T.M. performed the experiments and analyzed data. M.N. and A.S. wrote the manuscript with input from all authors. S.N. provided TCGA data and contributed to the interpretation of the results. M.J.M. provided critical feedback on the manuscript. All authors discussed the results and contributed to the final manuscript.

## Author declarations

The authors do not have anything to declare.

## Abbreviations

ATV: (Atorvastatin)
BafA1: (Bafilomycin A1)
CCCP: (Carbonyl cyanide m-chlorophenyl hydrazone)
DFP: (Deferiprone)
ER: (Endoplasmic Reticulum)
GKO: (GRAMD1C Knockout)
MBCD: (Methyl-β-Cyclodextrin)
PM: (Plasma Membrane)
Wt: (Wild-type)
ccRCC: (Clear Cell Renal Carcinoma)

Table I - GRAMD1C interactome

Proteins enriched from co-immunoprecipitation of GRAMD1C-EGFP (Table 1a) and GRAMD1C (ΔGRAM)-EGFP (Table 1b) were compared against proteins enriched from co-immunoprecipitation of EGFP tag alone. The log fold change (LogFC), P-values and protein IDs of the significant proteins are described.

Table II - The proteome of GRAMD1C depleted cells

Proteins identified from siGRAMD1C treated cells were compared against proteins from siNC treated cells. The log fold change (LogFC), P-values and protein IDs of the significant proteins are described.

