## Supplemental figures for "GRAMD1C regulates autophagy initiation and mitochondrial bioenergetics through ER-mitochondria cholesterol transport"

### Supplementary Figure 1

a.

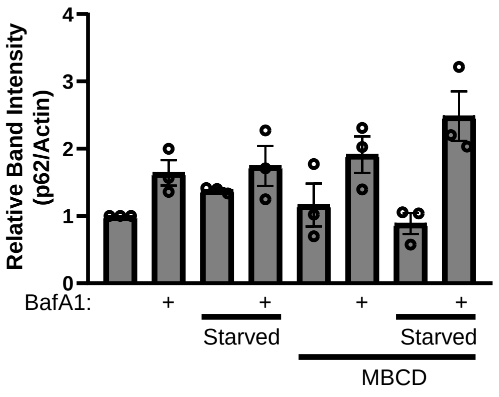


b.

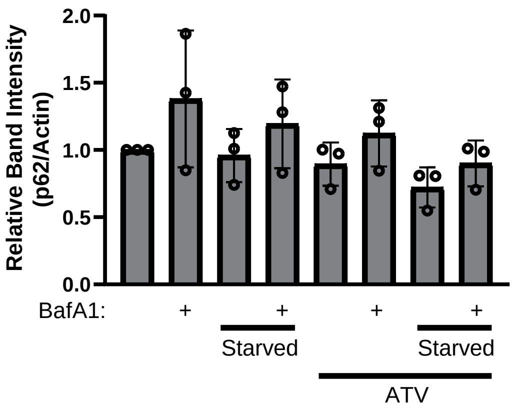


c.

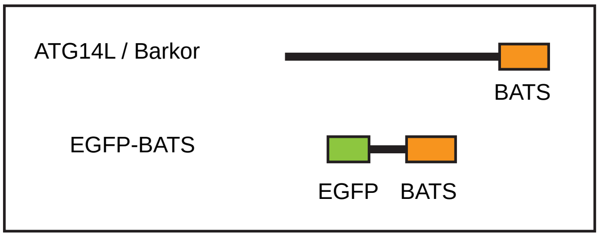


d.

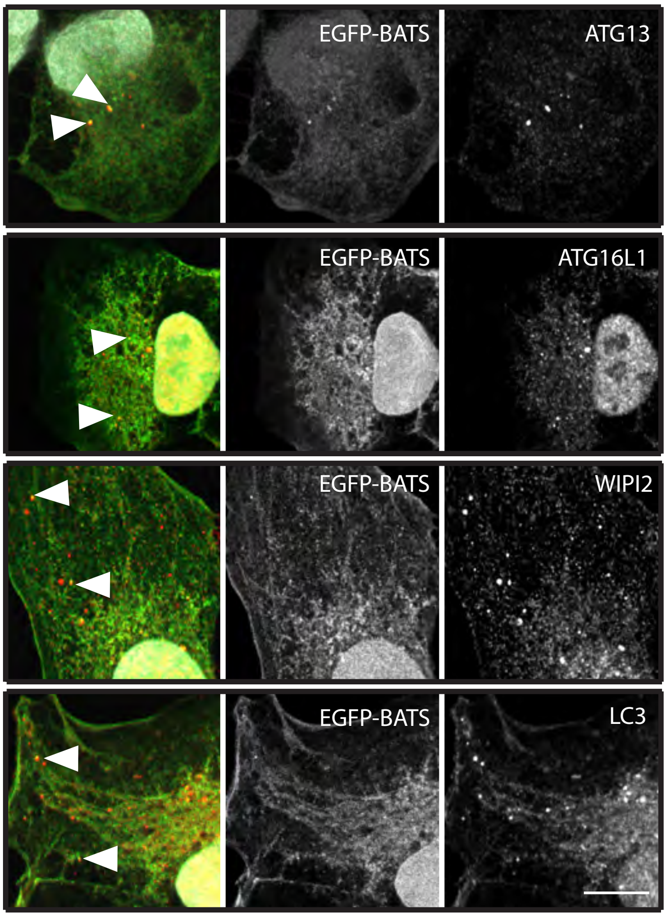

Supplementary figure 1 – EGFP-BATS are recruited to early autophagy structures

**a-b** The graphs represents the band intensities of p62 relative to actin and normalized to siNC from **a.** Figure 1a and **b.** Figure 1e. Error bars = SEM. **c** Graphical representation of ATG14L and its BATS domain, and the EGFP-BATS construct used. **d** U2OS cells stably expressing EGFP-BATS were starved in EBSS for 1 hr prior to fixation and immunostaining with the indicated antibodies prior to imaging with a confocal microscope. Scale bar = 10 µm.

### Supplementary Figure 2

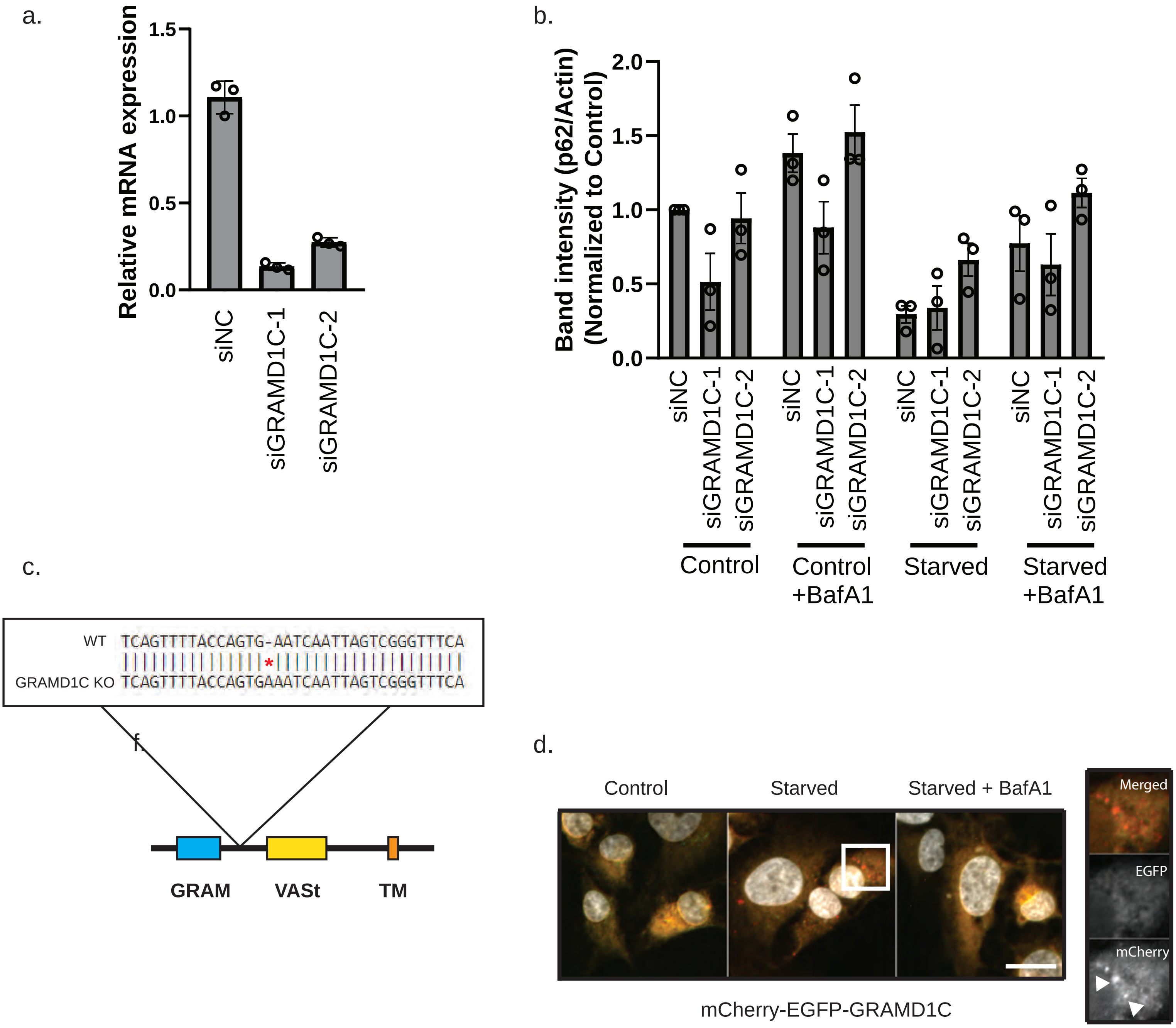

Supplementary figure 2 – GRAMD1C knockdown validation and degradation

**a** U2OS cells were treated with two different siRNAs against GRAMD1C for 72 hrs before mRNA extraction, cDNA synthesis and qPCR using primers against GRAMD1C. **b** U2OS cells were treated with siRNA against GRAMD1C for 72 hours prior to starvation in EBSS for 2 hrs ± BafA1. The graph represents the band intensities of p62 from Figure 3c. relative to actin and normalized to siNC. Error bar = SEM. **c** GRAMD1C Knockout Cells (GKO) KO were generated by CRISPR/Cas9, resulting in a single amino acid insertion in exon 7 of *GRAMD1C* leading to a premature stop codon (E245S*fs20). **d** U2OS cells stably expressing mCherry-EGFP-GRAMD1C were starved or not in EBSS (± BafA1) for 4 hrs, prior to fixation and widefield microscopy. Scale bar = 20 µm.

#### Supplementary Figure 3

a

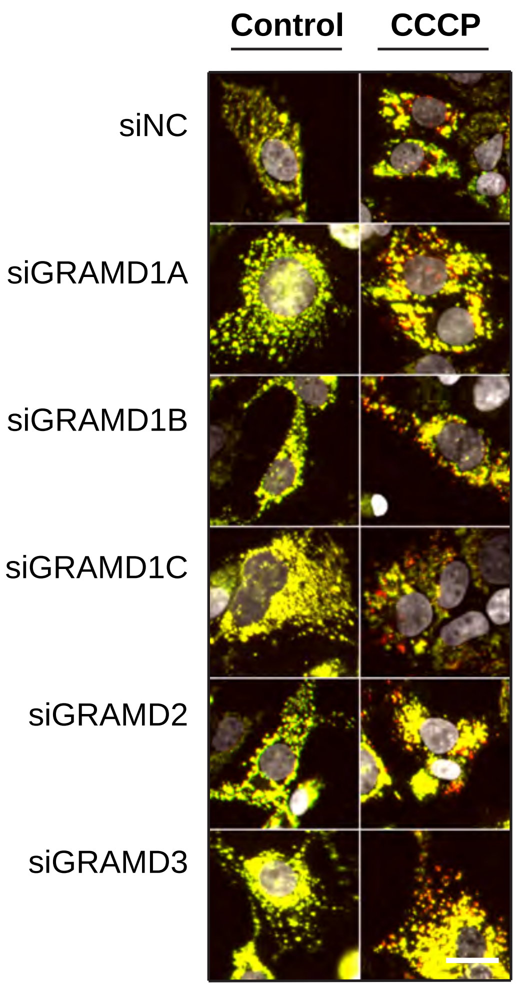


b

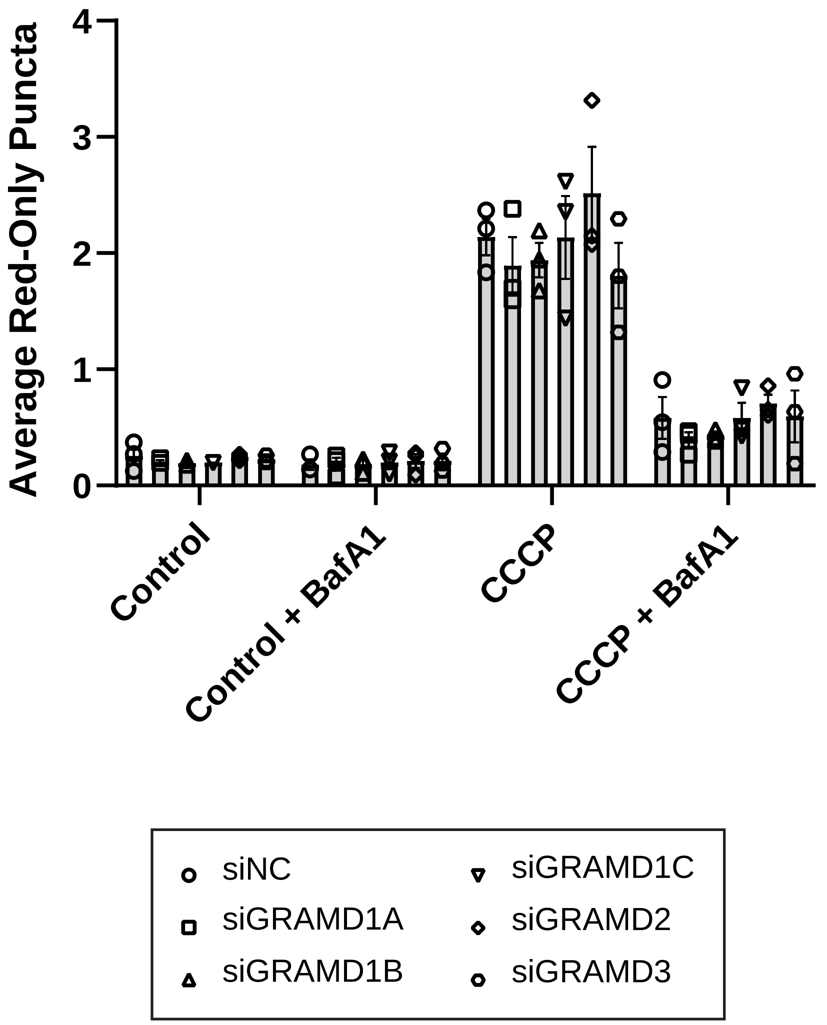


c

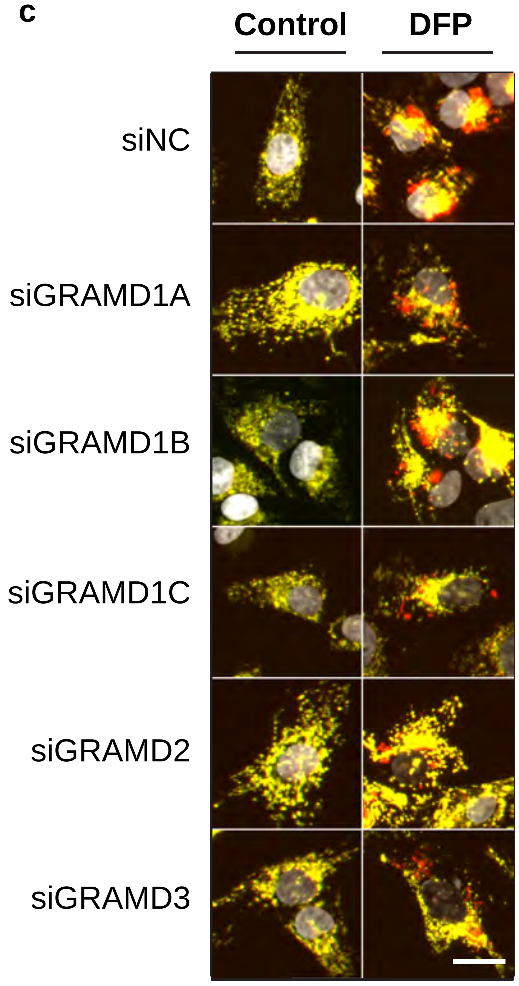


d

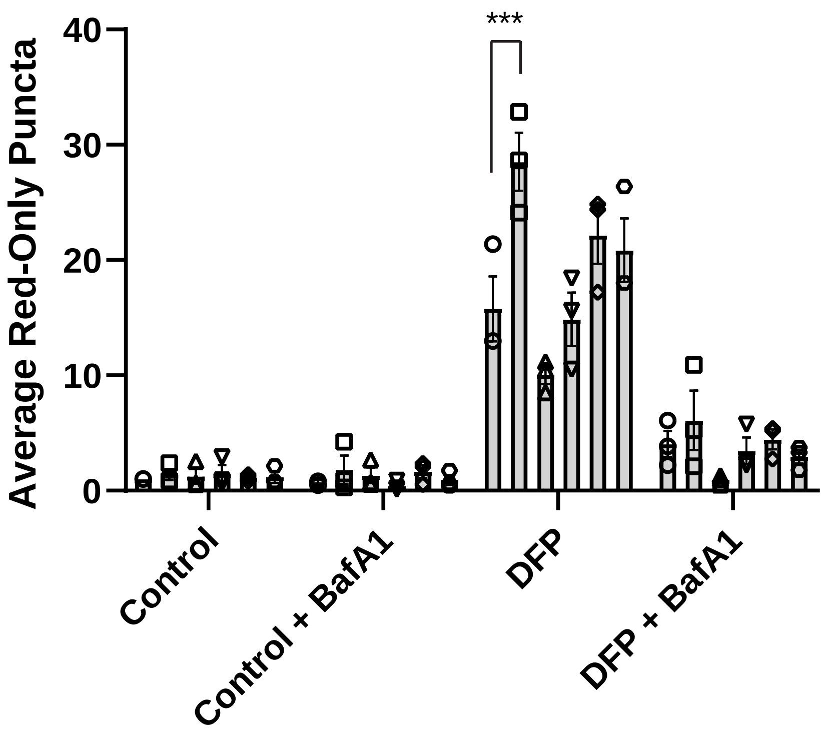

Supplementary figure 3 - GRAMD1C depletion on PARKIN-dependent/independent mitophagy

**a** U2OS cells stably expressing MLS (mitochondria localization signal)-mCherry-EGFP were treated with the indicated siRNA for 48 hrs before treatment with 1 mM DFP (+/- 100 nM BafA1) for 24 hrs. The cells were then fixed, and immediately imaged on a high throughput widefield microscope. Scale bar = 20 µM. **b** Quantification of images as shown in (**a**) from n = 3 experiments. The bars represent the average number of red only puncta per cell. Significance was determined using 2-way ANOVA followed by Tukey's comparison test. Error bar = SEM. **c** U2OS cells stably expressing MLS-mCherry-EGFP and untagged Parkin were treated with the indicated siRNA for 72 hrs before treatment with 10 µM CCCP (+/- 100 nM BafA1) for 8 hrs. The cells were then fixed, and immediately imaged on a high throughput widefield microscope. Scale bar = 20 µM. **d** Quantification of images as shown in c from n = 3 experiments. The bars represent the average number of red only puncta per cell. Significance was determined using 2-way ANOVA followed by Tukey's comparison test. Error bar = SEM. **** = $p<0.0001$, *** = $p<0.001$, ** = $p < 0.01$ and N.S. = not significant.

### Supplementary Figure 4

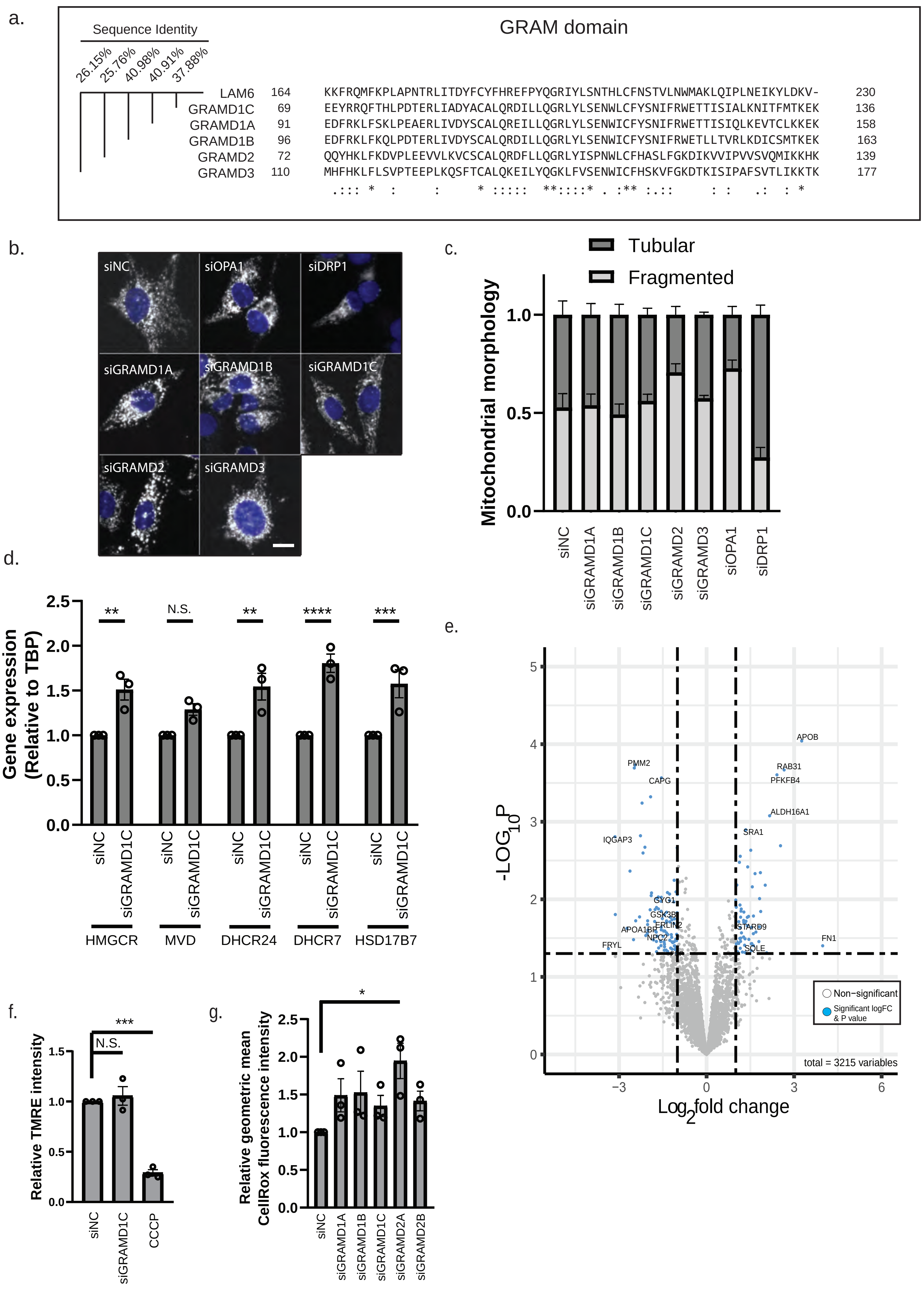

Supplementary figure 4 - GRAMD1C depletion leads to increased SREBP target gene expression

**a** GRAM domain sequence alignment of human *GRAMD1A*, *GRAMD1B*, *GRAMD1C*, *GRAMD2*, *GRAMD3* versus yeast *LAM6* using Clustal Omega. Sequence similarity values were obtained using BlastP. "." represents a conservation of weakly similar properties, ":" represents a conservation of strongly similar properties, "*" represents conserved amino acids. **b-c** Mitochondrial morphology was characterized in cells expressing MLS-mCherry-EGFP. The mitochondrial EGFP signal was analyzed with CellProfiler Analyst based on the mitochondrial area, texture, intensity distribution, intensity, and shape. Mitochondria from siDRP1 and siOPA1 treated cells were positive controls for tubular and fragmented mitochondria respectively. **d** U2OS cells were treated with siRNA against control (siNC) or GRAMD1C before RNA extraction and cDNA synthesis. The graph represents the gene expression of the indicated SREBP target genes relative to the expression of *TBP* and normalized to siNC. **e** Lysates from U2OS cells transfected control (siNC) or GRAMD1C siRNA were subjected to mass spectrometry analysis. The graph represents the difference in protein abundance in siGRAMD1C treated cells compared to siNC cells. Blue spots represent proteins that have $p < 0.05$ and a log fold change of > 1.3 or < -1.3. **f** U2OS cells treated with siRNA against control or GRAMD1C for 72 hrs were stained with TMRE and dye fluorescence intensity was measured by flow cytometry. CCCP treated cells were used as positive control. Significance was determined using 1-way ANOVA followed by Tukey's comparison test. Error bar = SEM. **g** U2OS cells were treated with the indicated siRNA for 72 hrs before staining with CellRox. The cells were then imaged using flow cytometry. Significance was determined using 1-way ANOVA followed by Tukey's comparison test. Error bar = SEM. **** = $p<0.0001$, *** = $p<0.001$, ** = $p < 0.01$ and N.S. = not significant.

### Supplementary Figure 5

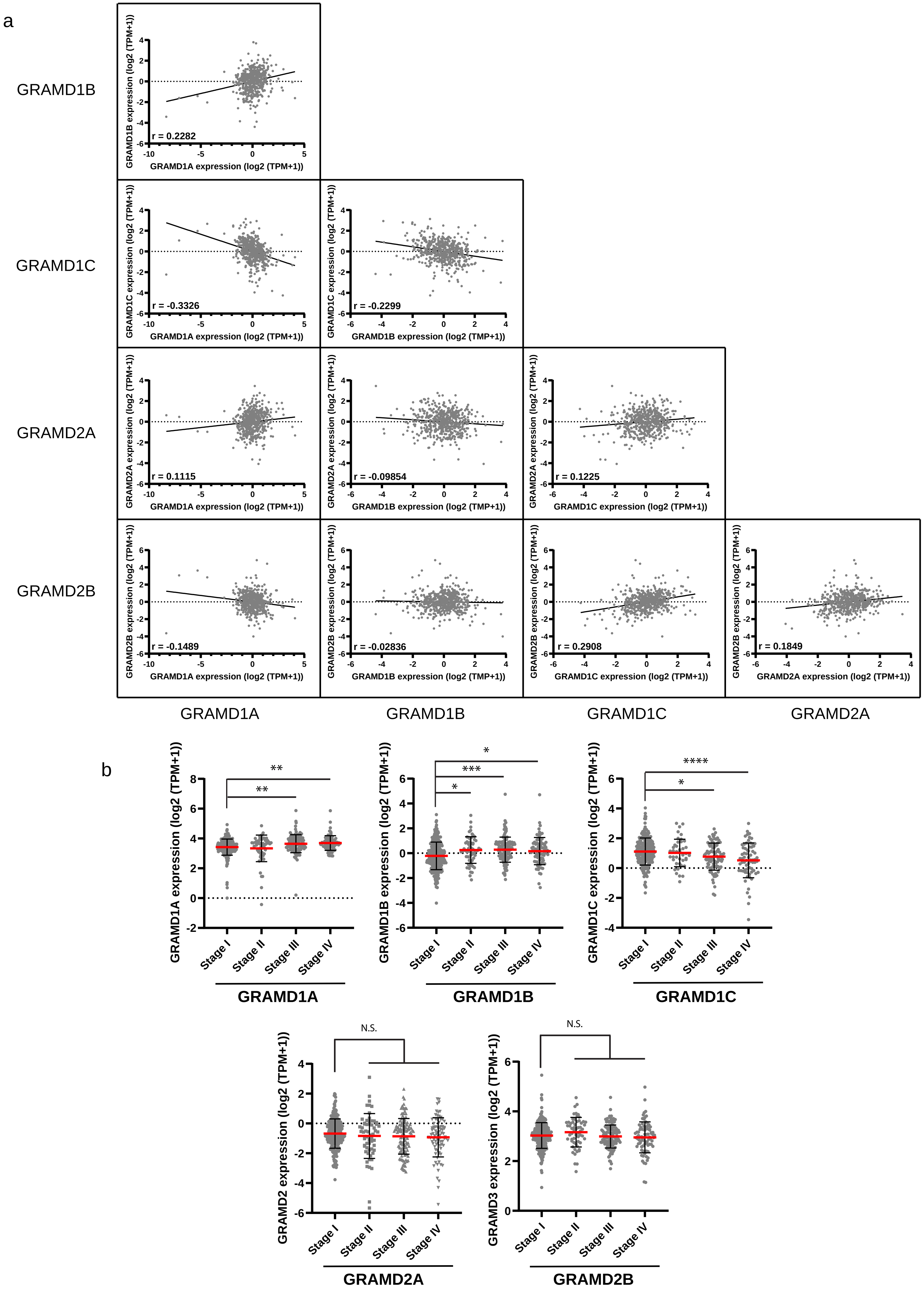

Supplementary figure 5 – The expression of GRAMs in ccRCC

**a** The expression of individual GRAM members from the TCGA-KIRC study were plotted against each other. The Pearson's correlation values are displayed in each graph. **b** The expression of individual GRAMs were plotted against cancer stage in the TCGA KIRC cohort. Significance was determined using 1-way ANOVA followed by Tukey's comparison test. Error bar = SEM. **** = $p<0.0001$, *** = $p<0.001$, ** = $p < 0.01$ and N.S. = not significant.

#### Supplementary Figure 6

a

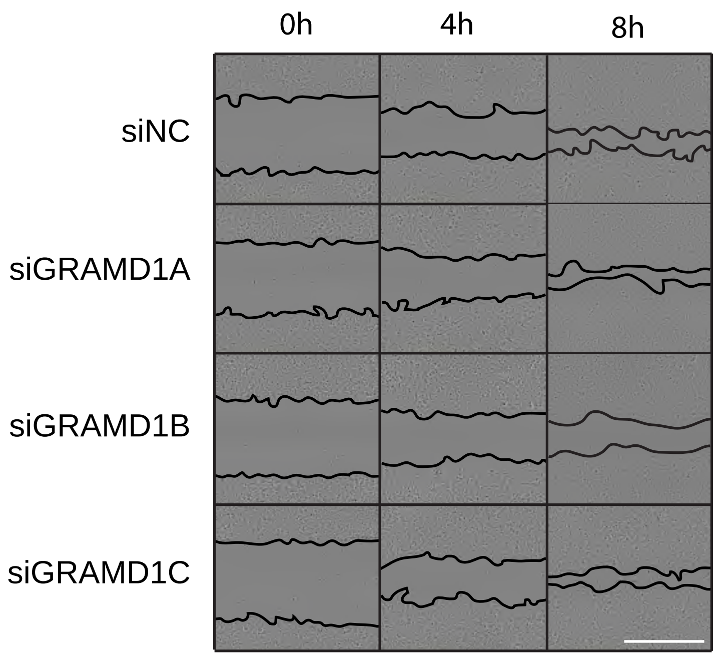


b

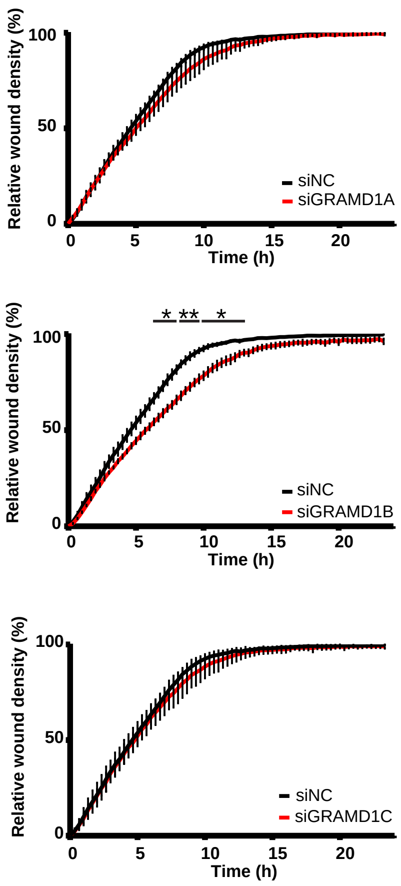


c

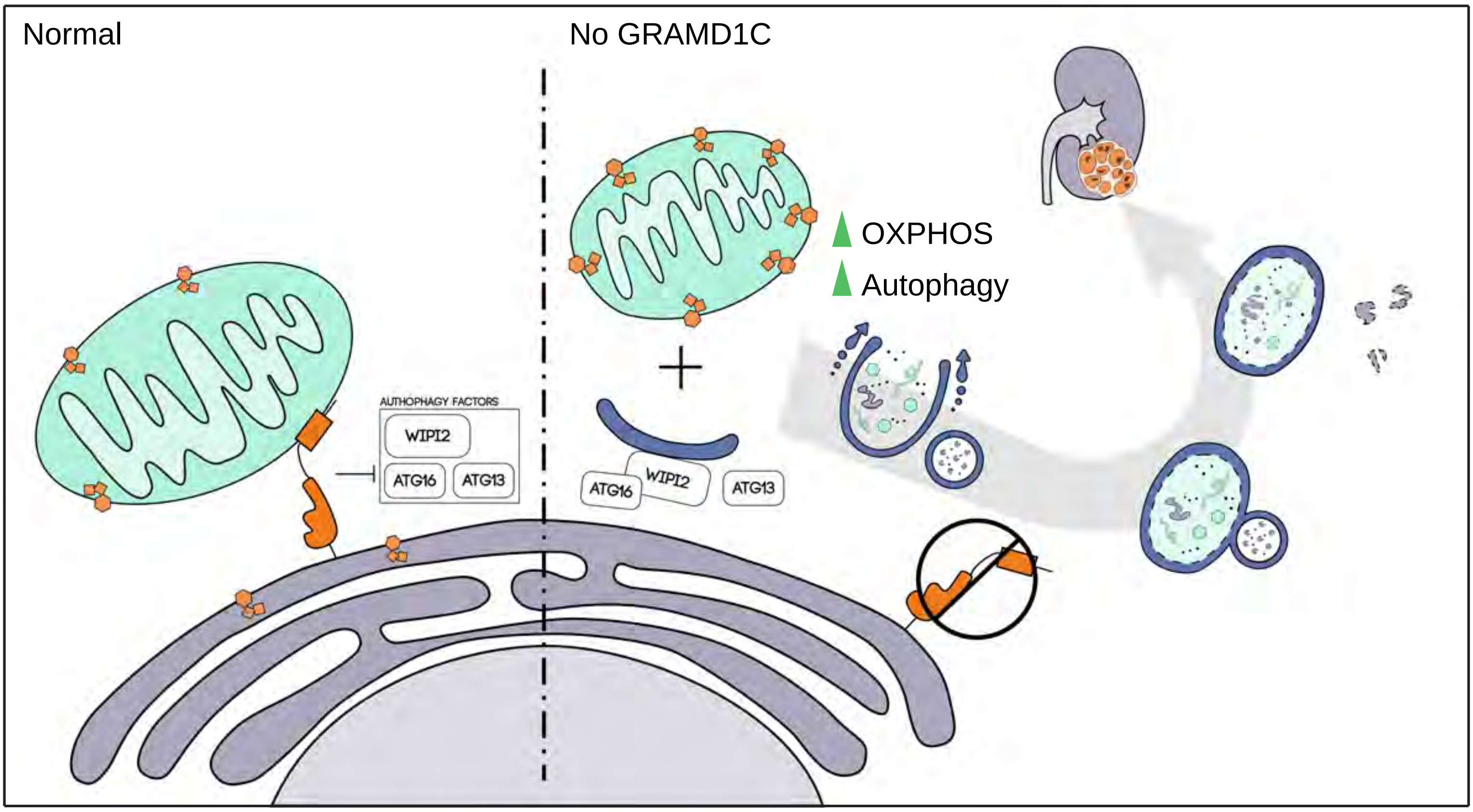

Supplementary figure 6 – GRAMD1B regulates cell migration in ccRCC cells

**a-b** 786-O cells treated with siRNA against the various GRAMD1s were grown to confluency. A rubber stopper was then used to generate a wound, and the cells were immediately imaged at 1 hr intervals using the Incucyte. **a** shows representative images taken at 0, 4 and 8 hrs. **b** The graph represents the relative wound density over time. Significance was determined using 2-way ANOVA from n = 3 individual experiments. Scale bar = 700 µm. ** = $p < 0.01$ and N.S. = not significant. **c** Graphical summary of the role of GRAMD1C as a negative regulator of autophagy. GRAMD1C localizes to the ER and interacts with mitochondria via its GRAM domain to facilitate cholesterol transport from mitochondria to the ER, thus limiting autophagosome biogenesis. Upon GRAMD1C depletion, the mitochondrial cholesterol abundance is increased, and we see increased membrane recruitment of early autophagic markers and increased autophagic flux, as well as increased mitochondrial respiration, thus contributing to ccRCC survival.
